# A generalizable Hi-C foundation model for chromatin architecture, single-cell and multi-omics analysis across species

**DOI:** 10.1101/2024.12.16.628821

**Authors:** Xiao Wang, Yuanyuan Zhang, Suhita Ray, Anupama Jha, Tangqi Fang, Shengqi Hang, Sergei Doulatov, William Stafford Noble, Sheng Wang

## Abstract

Nuclear DNA is organized into a compact three-dimensional (3D) structure that impacts critical cellular processes. High-throughput chromosome conformation capture (Hi-C) is the most widely used method for measuring 3D genome architecture, while linear epigenomic assays, such as ATAC-seq, DNase-seq, and ChIP-seq, are extensively employed to characterize epigenomic regulation. However, the integrative analysis of chromatin interactions and associated epigenomic regulation remains challenging due to the pairwise nature of Hi-C data, mismatched resolution between Hi-C and epigenomic assays, and inconsistencies among analysis tools. Here we propose HiCFoundation, a Hi-C-based foundation model for integrative analysis linking chromatin structure to downstream regulatory function. HiCFoundation is trained from hundreds of Hi-C assays encompassing 118 million contact matrix submatrices. The model achieves state-of-the-art performance on multiple types of 3D genome analysis, including reproducibility analysis, resolution enhancement, and loop detection. We further demonstrate the model’s generalizability through genome architecture analysis of 316 species. Notably, by enhancing low-coverage experimental Hi-C data, HiCFoundation reveals genome-wide loop loss during differentiation of hematopoietic stem and progenitor cells (HSPCs) to neutrophils. Additionally, HiCFoundation is able to predict multiple types of epigenomic activity from Hi-C input and further interprets the link between Hi-C input and epigenomic output to reveal the relationship between chromatin conformation and genome function. Finally, HiCFoundation can analyze single-cell Hi-C data, shedding light on genome structure at single-cell resolution. HiCFoundation thus provides a unified, e”cient, generalizable, and interpretable foundation for genome architecture, single-cell and multi-omics analysis across species, paving the path for systematically studying genome 3D architecture and its regulatory mechanisms.

## Main

Genomes are physical objects organized in a compact 3D structure within the cell nucleus. This spatial organization impacts various genome functions, such as DNA replication, transcription, cell differentiation, and cell senescence, in ways that cannot be fully understood only from the one-dimensional DNA sequence [1]. As a result, many experimental methods have been developed over the last two decades to measure the 3D conformation of DNA within the nucleus. In particular, high-throughput chromosome conformation capture (Hi-C) [2] provides a genome-wide “contact matrix,” where the two axes are genomic loci and values in the matrix represent proximity in 3D. Single-cell Hi-C (scHi-C) [3] further extends Hi-C by measuring genome architecture in individual cells. In parallel, epigenomic assays such as ATAC-seq [4], DNase-seq [5], transcription factor (TF) ChIP-seq [6], and histone ChIP-seq [7] measure, respectively, chromatin accessibility, transcription factor binding, and histone post-translational modifications. Because Hi-C and these epigenomic assays provide complementary information about the genome, their joint analysis can deepen our understanding of the links between genome architecture and epigenomic regulation.

However, two key challenges hinder progress towards jointly analyzing these data. First, Hi-C data and epigenomic assays typically have different resolutions. Epigenomic assays are usually represented at basepair resolution, whereas Hi-C data is often analyzed at 10 kb or coarser resolution due to the limitation of sequencing depth; e.g., doubling the resolution of the Hi-C contact matrix requires a fourfold increase in sequencing cost. This lack of depth makes detecting some 3D structures especially challenging. Notably, the robust detection of chromatin loops, a dynamic 3D genome feature that is closely associated with gene regulation [8], requires deeply sequenced Hi-C data. In many settings, Hi-C resolution is further constrained by the limited availability of biologically important cell types, such as hematopoietic stem and progenitor cells (HSPCs) and rare populations of immune cells. The scHi-C assay partially addresses this issue by providing cell-type-specific measurements of chromatin architectures. However, the resolution mismatch becomes even more severe in the scHi-C assay, where its coverage is orders of magnitude lower than bulk Hi-C. The second challenge is that different types of 3D structures exhibit complex interdependencies, but separate tools are typically used to identify these structures, which can yield inconsistent results. For example, a common feature of 3D chromatin arises due to cohesin-mediated loop extrusion [9]. As a result, such loops typically link together the two boundaries of topologically associating domains (TADs), so calling TADs and loops independently is potentially problematic. Therefore, there is a pressing need for a unified computational framework that leverages the dependencies among 3D structures and epigenomic regulation and facilitates the analysis of Hi-C data, scHi-C data and epigenomic data for systematically studying genome architecture and its functional implications.

This need for a unified framework aligns perfectly with the foundation modeling paradigm in machine learning. The key idea behind a foundation model is to simultaneously tackle many tasks by learning a shared model using a large collection of datasets [10]. In particular, foundation models are typically developed using a two-stage approach. First, in the pre-training stage, a deep neural network model is trained in a self-supervised fashion on massive quantities of unlabeled data. Second, this foundation model is separately fine-tuned in a supervised fashion for each of the downstream tasks using a smaller amount of task-specific labeled data. Intuitively, a foundation model that learns from a large number of Hi-C experiments can increase the signal-to-noise ratio of its input data by learning patterns from many contact matrices, leveraging dependencies among different Hi-C applications, and later enabling joint analysis with epigenomic assays in a data-driven way. Recognizing the transformative potential of foundation models, the wealth of genomic information embedded in Hi-C data, and the intrinsic similarity between Hi-C and scHi-C, we hypothesized that a foundation model approach could provide a unified framework for solving diverse genome architecture-related problems, including analysis of bulk and single-cell data as well as translation between Hi-C and epigenomic data.

To this end, we developed HiCFoundation, a foundation model for genome architecture and function analysis. HiCFoundation is trained from hundreds of Hi-C assays in 81 human cell lines or tissue types (“biosamples”), encompassing 118 million contact submatrices. We propose an encoder-decoder pre-training architecture that exploits a novel patch-wise contrastive loss to overcome the extreme sparsity in Hi-C data. The trained HiCFoundation model is able to generate three levels of general-purpose embeddings: embeddings of spatial contact information for a full chromosome, for a set of contiguous genomic loci (i.e., a submatrix of the Hi-C contact matrix), and for a single genomic locus (i.e., one row or column of the Hi-C contact matrix). These three embeddings enable diverse types of Hi-C analyses, including investigating genome architecture at different scales and resolutions.

We evaluate the multi-scale embeddings provided by HiCFoundation on many different tasks. We demonstrate that HiCFoundation offers a generalizable analysis engine for genome architecture, delivering state-of-the-art performance across various tasks, including reproducibility analysis, resolution enhancement, and chromatin loop detection. We further show the cross-species generalizability of HiCFoundation by empirically validating the model using Hi-C data from 316 species, many of which exhibit substantial evolutionary divergence from humans. HiCFoundation also reveals a genome-wide loss of chromatin loops during differentiation of HSPCs to neutrophils, while loops around genes essential for neutrophil function are maintained and refined for optimal expression. Additionally, loss of nuclear lamin B1 reduces the overall loop strength in both HSPC and neutrophils. Furthermore, HiCFoundation extends beyond genome architecture analysis by predicting epigenomic activity, such as chromatin accessibility (DNase-seq/ATAC-seq), transcription factor binding (TF ChIP-seq), and histone modifications profiles (histone ChIP-seq), directly from Hi-C data. To our knowledge, this is the first model that successfully infers genome activity from the coarse genomic contact maps provided by Hi-C. Our interpretability analysis using HiCFoundation [11, 12] further reveals the relationship between chromatin loop structure and corresponding epigenomic features, thereby enabling systematic hypothesis generation at the interface between genome architecture and function. Finally, HiCFoundation alleviates the sparsity challenge in scHi-C data by leveraging patterns from bulk Hi-C to enhance scHi-C contact matrices. Compared with state-of-the-art methods, our model achieves superior performance across four scHi-C datasets and reveals genome compartments that are validated by ATAC-seq data. HiCFoundation thus offers a unified foundation for jointly analyzing genome architecture and associated epigenomic activity.

## Results

### HiCFoundation is pre-trained from unlabeled Hi-C experiments

The key idea of HiCFoundation is to train one model from a large number of Hi-C experiments and fine-tune it to quickly adapt to new cell types, species and modalities (**Fig. 1**). To this end, HiCFoundation leverages 1015 Hi-C experiments from over 300 species, sourced from the ENCODE [13], 4D Nucleome (4DN) [14], and DNA Zoo [15] datasets. Pre-training is carried out in a self-supervised fashion on Hi-C data from a designated set of training chromosomes, across 101 human biosamples (**Fig. 1a, Supplementary Table 1**). Subsequently, the model is fine-tuned using task-specific signals for different downstream analyses in a supervised fashion. Following training, HiCFoundation is benchmarked on 117 Hi-C experiments from 18 human biosamples, as well as 494 Hi-C experiments from more than 300 species.

**Figure 1:**
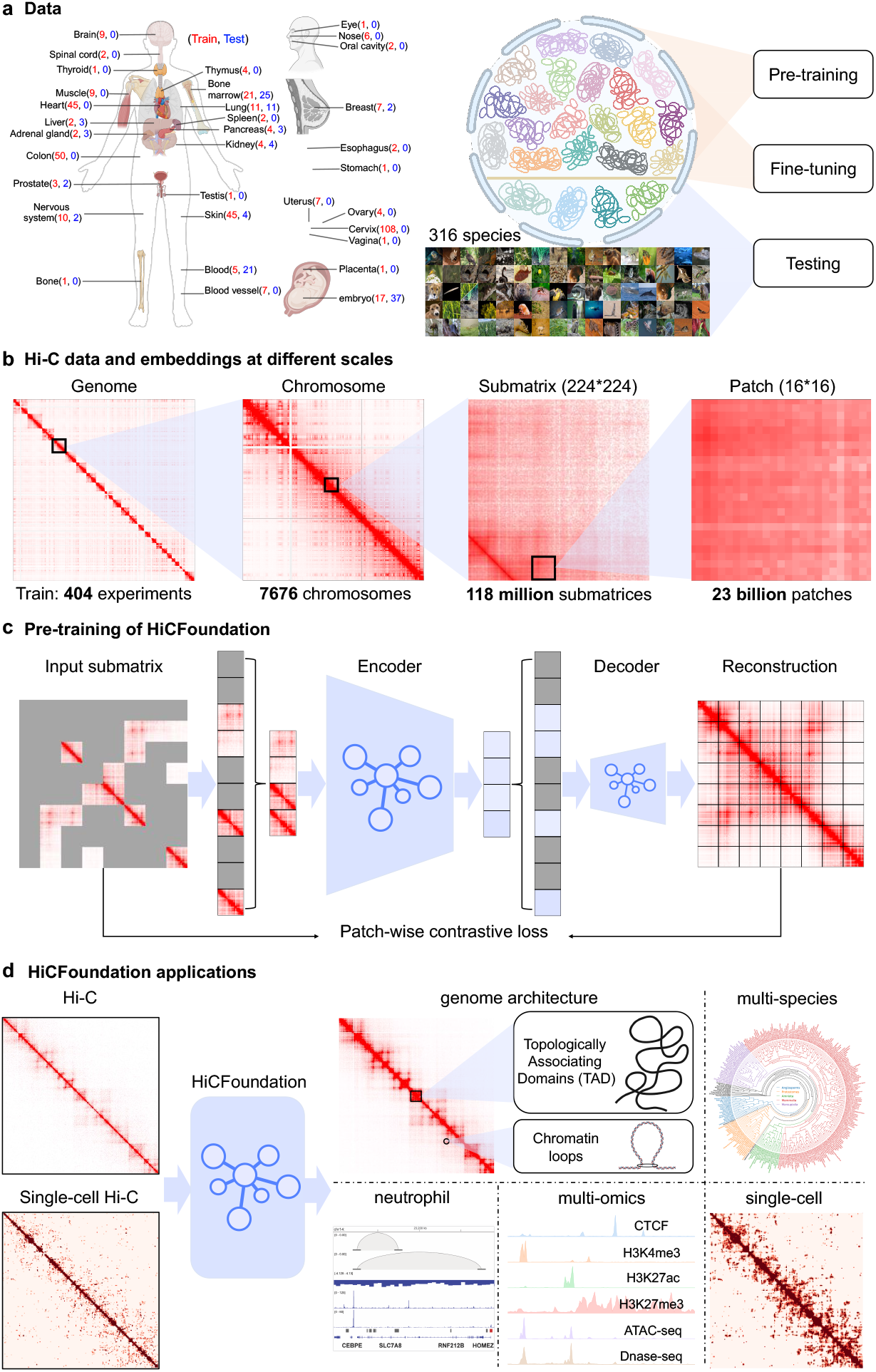
Overview of HiCFoundation. **a**. Illustration of the data used in the development of HiCFoundation. Human Hi-C experiments are used for training and testing of HiCFoundation, including 521 Hi-C experiments spanning 119 cell lines from 33 organs. Human chromosomes are split into training and testing chromosomes, where pre-training and fine-tuning are only done using the training chromosomes. HiCFoundation is subsequently benchmarked on testing chromosomes from human cell lines and all chromosomes from 316 different species. **b**. Illustration of Hi-C data and embeddings at different scales. We collected 404 Hi-C experiments, encompassing a total of 7,676 training chromosomes. From these, we extracted 118 million non-overlapping submatrices, each sized 224 ×224, as inputs for the HiCFoundation model. These submatrices were further divided into 23 billion non-overlapping patches, serving as inputs for the model. HiCFoundation is designed to generate embeddings at different scales corresponding to these data scales. **c**. Pre-training framework of HiCFoundation. HiCFoundation is pre-trained using masked Hi-C submatrices as input, optimizing for the reconstruction of the full submatrix. Specifically, its encoder processes visible patches to generate their embeddings, while the decoder leverages these embeddings along with masked token embeddings to reconstruct all patches. This framework is trained using a novel patch-wise contrastive loss and SSIM loss. **d**. HiCFoundation applications. Following pre-training, HiCFoundation is fine-tuned and tested for diverse downstream tasks, including genome architecture, multi-species, neutrophil, multi-omics and single-cell analyses.

During the pre-training stage, we first randomly sampled 118 million submatrices from the training set **(Fig. 1b)**. Each Hi-C submatrix is further divided into regular, non-overlapping patches of 16 ×16 bins, which serve as the input for the deep learning model (**Fig. 1b**). During the pre-training stage, HiCFoundation takes partially masked Hi-C submatrices as input and is optimized to reconstruct full submatrices (**Fig. 1c**). Inspired by masked autoencoders [16], this reconstruction is implemented using an encoder-decoder architecture based on the Vision Transformer [17, 18]. We introduced a novel patch-wise contrastive loss term to alleviate the sparsity within certain patches, especially those that are far away from the diagonal [19] (see **Methods**). During the fine-tuning stage, the encoder is frozen and shared across various tasks, while the decoder is optimized to be task-specific. As a result, we use a larger encoder (304M parameters) and a smaller decoder (26M parameters) to maximize learning capacity during pre-training while minimizing the fine-tuning cost. Pre-training required two weeks on a server with 8 A100 (80GB) GPUs.

Before applying HiCFoundation to downstream applications, we first examined the reconstruction of submatrices in the test set to ensure the success of the pre-training. On this task, we compared HiCFoundation with three baselines: using the masked Hi-C as a prediction (“raw”), filling masked patches with the unmasked diagonal average values (“diag avg”), and filling masked patches with the mean values of the unmasked submatrix regions (“submat avg”). HiCFoundation achieved the best performance, with 17.7% structural similarity index measure (SSIM) improvement, 15.2% Pearson correlation improvement, and 19.1% Spearman correlation improvement over the best-performing baseline (**Supplementary Table 2, Extended Data S1a**). Visualizations of reconstructed examples across diverse species from the test set are shown in **Extended Data S1b**. The exceptional performance of HiCFoundation on the reconstruction task validates the success of its pre-training procedure, further motivating us to evaluate its performance on real-world applications.

After pre-training, HiCFoundation is fine-tuned and evaluated for Hi-C analysis to investigate genome architecture across species, differential Hi-C analysis to study chromatin organization changes in neutrophil, multi-omics analysis to explore the relationship between genome architecture and function, and single-cell analysis to characterize the structural organization of individual cells (**Fig. 1d**).

### HiCFoundation enables multiple types of chromatin architecture analysis

Hi-C has emerged as an indispensable tool for characterizing 3D chromatin architecture, supported by a wide array of computational methods [20, 21, 22, 23, 24, 25, 26, 8]. To validate HiCFoundation’s capacity for chromatin architecture analysis, we focus on three important tasks.

- **Reproducibility analysis** evaluates the consistency between biological replicates and is critical for Hi-C quality control. Reproducibility measures are important for deciding whether two replicates can be pooled. It is also an essential step for differential chromatin architecture analysis, where the differences between various conditions can only be trusted if they exceed the differences between biological replicates. To this end, several reproducibility approaches have been developed and widely adopted [23, 27, 28, 29, 30].
- **Chromatin loop detection** identifies pairs of genomic loci that are in close spatial proximity relative to their local background [8, 31, 32]. Loop detection can provide insights into regulatory interactions and genome organization. For example, some loops are mediated by cohesin-driven loop extrusion, which is closely associated with gene regulation [9].
- **Resolution enhancement** aims to increase the effective sequencing depth of Hi-C data, thereby allowing for the detection of finer-scale chromatin features. Given that experimental doubling of resolution incurs a fourfold increase in cost, numerous computational methods for resolution enhancement have been developed to complement experimental approaches [33, 34, 35, 21, 36, 20, 37].

For each of these tasks, HiCFoundation takes Hi-C data as input, processed by the pre-trained encoder and a task-specific decoder. The model outputs a submatrix embedding for reproducibility analysis, a list of detected chromatin loops for loop detection, or an enhanced Hi-C matrix for resolution enhancement (**Fig. 2a**). The encoder is frozen and shared across tasks, whereas the decoder is fine-tuned in a task-specific fashion (**Extended Data S2**, see details in **Methods**).

**Figure 2:**
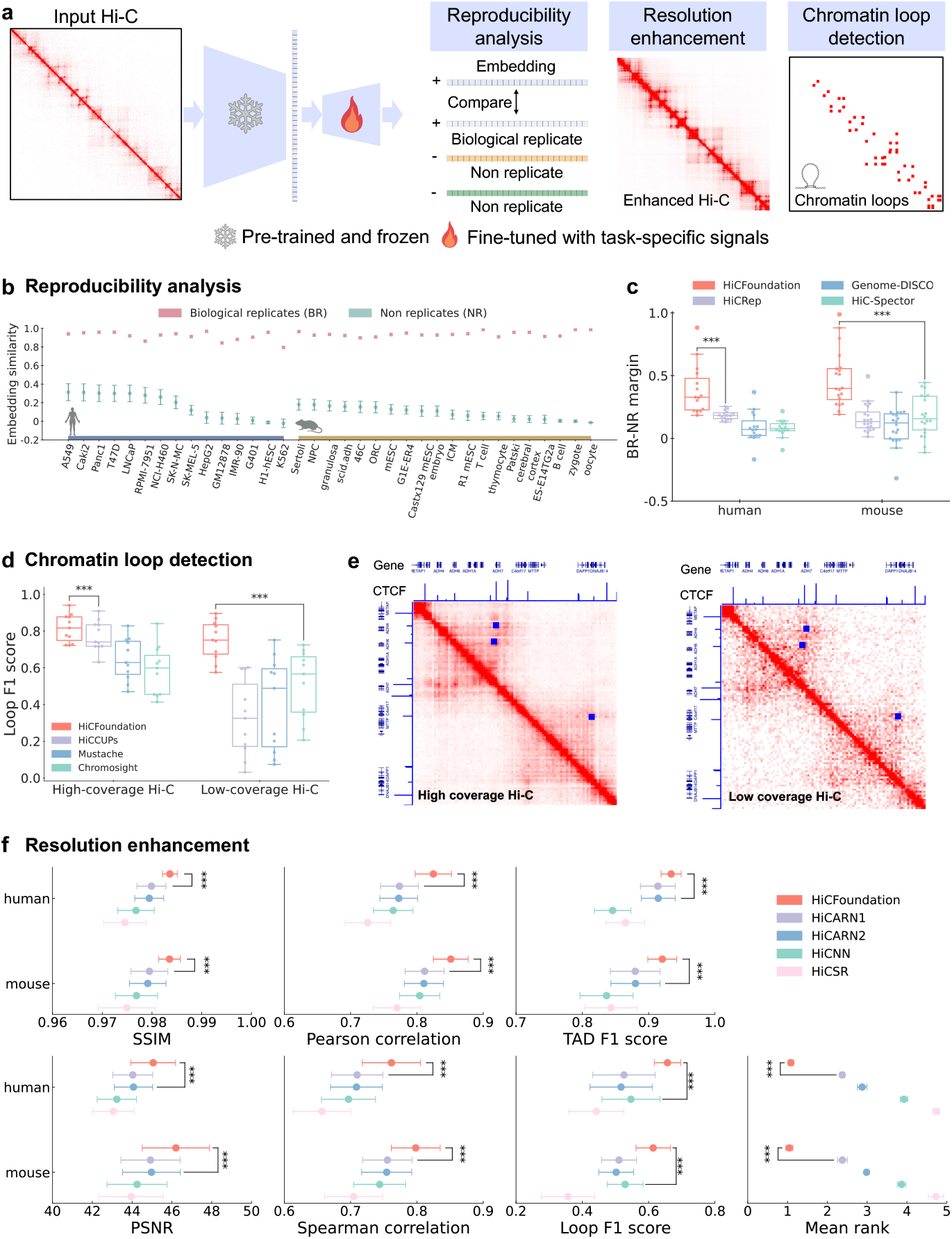
HiCFoundation for chromatin architecture analysis. **a**. Overview of HiCFoundation for different chromatin architecture analyses from Hi-C. The model takes a Hi-C submatrix as input, processed by a frozen encoder and a task-specific decoder, and outputs an embedding vector for reproducibility analysis, an enhanced Hi-C matrix, or a list of detected chromatin loops. **b**. Evaluation on reproducibility analysis. Box plot comparing pairwise embedding similarities between biological replicates (BR) and non-replicates (NR) across various biosamples from human and mouse. The embedding is derived using HiCFoundation. **c**. Box plot comparing BR–NR margin across different methods. The BR–NR distribution of other methods across cell lines are included in **Extended Data S3. d**. Box plot comparing the loop detection F1 scores across different methods on high-coverage and low-coverage Hi-C. **e**. Case studies of the detected loops from high-coverage and low-coverage Hi-C on the IMR-90 cell line (ENCSR852KQC) from the ADH gene family region. Hi-C data, corresponding CTCF ChIP-seq signals (ENCSR000EFI), reference genes, and loop detections (blue squares) are also included. **f**. Box plots comparing HiCFoundation with competing methods on resolution enhancement using seven different evaluation measures: SSIM, PSNR, Pearson correlation, Spearman correlation, TAD F1 score, Loop F1 score, and mean rank. Mean rank is the mean of the rank in each of the other six metrics. The datasets used for reproducibility analysis, loop detection, and resolution enhancement are detailed in **Supplementary Tables 3, 5, and 7**, respectively. The detailed scores of these metrics are provided in **Supplementary Tables 4, 6, and 8**, respectively. “***” indicates significance based on the sign test with *p <* 10−^3^.

For reproducibility analysis, HiCFoundation derives submatrix embeddings to calculate a similarity score between two Hi-C matrices. A good Hi-C similarity metric should assign a higher score to pairs of Hi-C experiments that are derived from the same or similar biological samples (“biological replicates”—BR) than to pairs derived from different samples (“non-replicates”—NR) [38, 30]. Accordingly, HiCFoundation takes three input submatrices from BR1, BR2, and NR and generates three corresponding embeddings (**Extended Data S2a**, see **Methods**). We fine-tune the lightweight decoder using a triplet margin loss by minimizing the distance between BR1 and BR2 and maximizing the distance between BR1/BR2 and NR.

The learned embeddings of the fine-tuned model, HiCFoundation-Repro, yield consistently higher similarity between BRs than NRs (**Fig. 2b** and **Extended Data S3**) across various test set biosamples from human and mouse, demonstrating the model’s ability to effectively distinguish between BR and NR pairs. We observed that HiCFoundation achieves the largest BR–NR margin, which measures the difference between the score of the BR pair and the 95th percentile of reproducibility scores of NR pairs within each biosample, among all methods, with 110% and 158% improvement relative to the second-best method on the human and mouse test sets, respectively (**Fig. 2c, Supplementary Table 4**).

Next, we examined whether HiCFoundation can provide robust loop detection across varying coverage levels. To achieve this, we fine-tuned two HiCFoundation-Loop models for high-coverage (HC) and low-coverage (LC) Hi-C data, respectively. In each case, we train and evaluate the model using consensus loop calls produced by HiCCUPs [8] on two replicated HC Hi-C experiments at 10 kb resolution (see **Methods**). Following pixel-level predictions by HiCFoundation-Loop, the mean-shift algorithm [39] is employed to cluster pixel-wise predictions to yield a final set of loop calls. Comparing the loop F1 score of different methods on various biosamples in the test set (**Fig. 2d, Supplementary Table 6**), HiCFoundation-loop achieves mean loop F1 scores of 81.6% and 75.1% in the HC and LC settings, substantially outperforming other methods (sign test *p <* 0.001 relative to second best HiCCUPs on HC Hi-C data, and second best Chromosight on LC Hi-C data). Loops detected in the ADH gene family region [40] in IMR-90 data exhibit strong consistency with the corresponding CTCF ChIP signals and gene annotations (**Fig. 2e**). We successfully captured 100% of the target loops (consensus loops in **Extended Data S4a** by HiCCUPs on two biological replicates). In contrast, HiCCUPs failed to detect any loops in this region from the low-coverage Hi-C (**Extended Data S4b**). These results indicate HiCFoundation-Loop’s ability to detect chromatin loop architecture accurately at varying coverage levels.

Finally, we evaluated HiCFoundation on resolution enhancement, where the model takes as input an LC Hi-C submatrix and is optimized to generate the corresponding HC submatrix. The input and output are both raw count matrices. We intentionally deviated from the common practice of using normalized Hi-C for this task because we identified a systematic problem associated with the Hi-C normalization approach adopted by previous methods (**Supplementary Note 1**). The fine-tuned model, HiCFoundation-Reso, was then compared to HiCARN [33], HiCNN [34] and HiCSR [41], each of which was re-trained using the same data and chromosome split to ensure a fair comparison. For evaluation, we adopt the six performance measures outlined in previous work [42]: SSIM, peak signal-to-noise ratio (PSNR), Pearson correlation, Spearman correlation, TAD F1 score, and loop F1 score (with loops called by HiCFoundation-Loop). Among all six different evaluation metrics, HiCFoundation achieves the best performance across both the mouse and human test sets (**Fig. 2f, Supplementary Table 8**). To summarize across all six performance measures, we defined a mean rank metric, which is the average rank of a given method relative to its competitors across all six metrics. HiCFoundation achieves a mean rank of 1.08 on human and 1.05 on mouse, which is substantially lower than that of the next best method (2.38/2.38 for HiCARN1, **Fig. 2f**), demonstrating the superior performance of HiCFoundation on resolution enhancement.

To assess the necessity of pre-training, we also include a “No pretrain” baseline, which uses the same network architecture but with random initialization, where the entire backbone is optimized using only task-specific signals. Across three chromatin architecture analyses, HiCFoundation consistently outperforms the No-pretrain model (**Extended Data S5**). The substantial performance gains achieved by HiCFoundation in this setting highlight the effectiveness of pre-training on massive unlabeled Hi-C data. Additionally, HiCFoundation is substantially more e”cient in downstream analyses, achieving approximately 10 times faster training than the No-pretrain model by requiring fine-tuning on only a small decoder rather than the entire network.

To further assess the utility of HiCFoundation, we benchmarked it on the DNA Zoo dataset [15], which contains Hi-C data from more than 300 primarily mammalian species. Given that HiCFoundation has been pre-trained and fine-tuned solely on human data, the DNA Zoo data allows us to test the model’s ability to generalize across species. Using the same settings and evaluation protocols as before, we compared HiCFoundation-Reso against other methods on the resolution enhancement task. Across all our performance measures, HiCFoundation-Reso again consistently outperforms its competitors (**Fig. 3a, Supplementary Table 10**). This result indicates the applicability of HiCFoundation across a wide variety of species. To test HiCFoundation’s generalizability, we further compared HiCFoundation against No-pretrain under different coverage settings by varying the downsampling ratio in our evaluation framework. HiCFoundation consistently delivers better performance relative to the No-pretrain model (**Fig. 3b**). Most notably, we observed that the improvement of HiCFoundation over No-pretrain is higher in non-mammalian species than mammalian species across different evaluation metrics (**Fig. 3c and Extended Data S6**), suggesting that the pre-training stage enhances the generalizability of HiCFoundation. The self-supervised training during pre-training eliminates inductive bias, allowing the model to better adapt to evolutionarily distant species.

**Figure 3:**
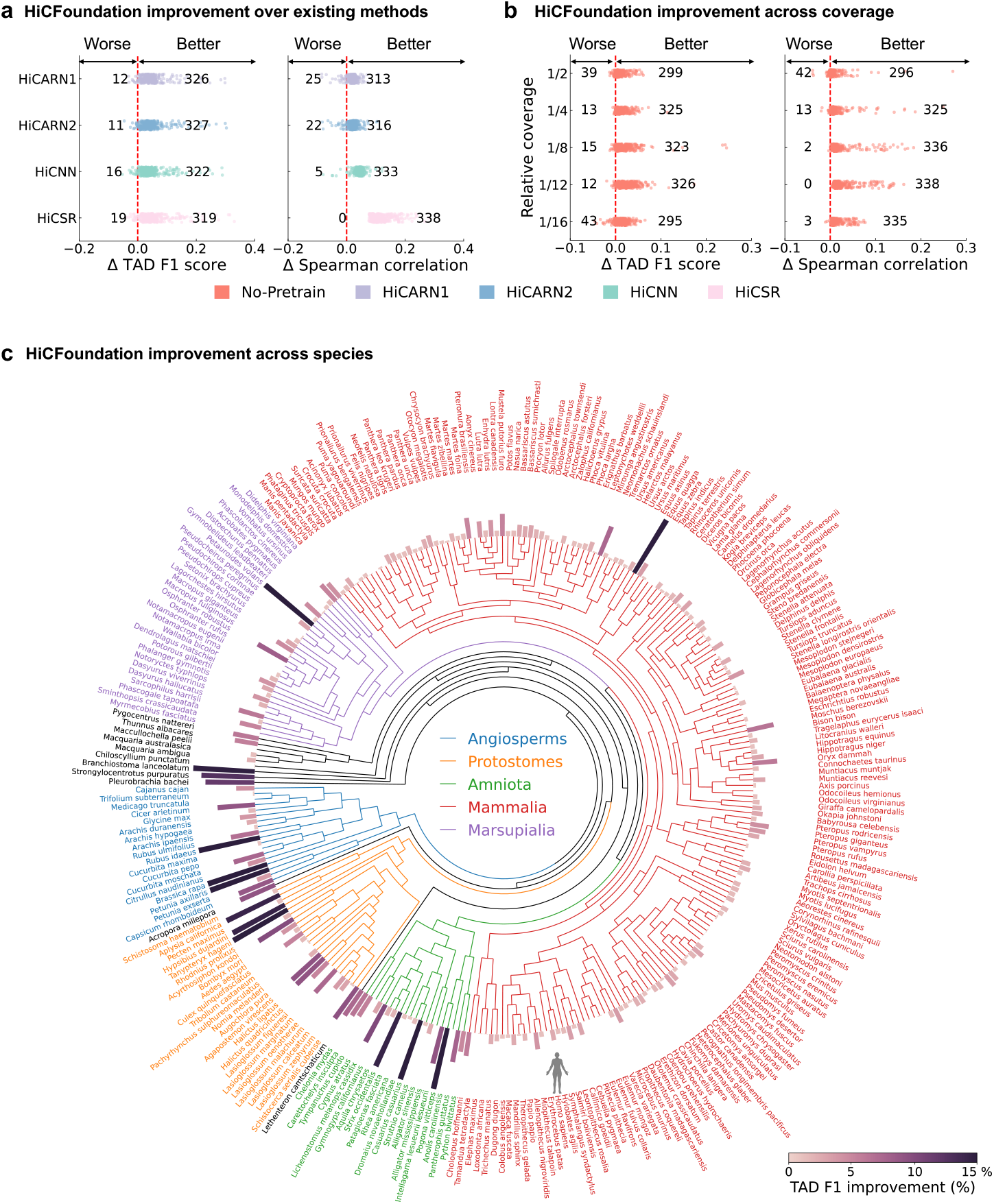
Multi-species evaluation of HiCFoundation on DNA Zoo dataset. **a**.Dot plots showing the improvement of HiCFoundation over existing methods on resolution enhancement. Each dot shows the difference between HiCFoundation and other methods for a Hi-C experiment from a specific species in terms of the metric shown on the x-axis. A positive (negative) difference indicates HiCFoundation is better (worse) than the competing method. The numbers of differences greater than or less than zero are also shown in the figure. **b**. The improvement of HiCFoundation over No-pretrain across different coverages relative to the depth of the raw data. **c**. TAD F1 score improvement of HiCFoundation over No-pretrain across 316 species. Results using other evaluation metrics are available in **Supplementary Table 10 and Extended Data S6**.

### HiCFoundation reveals 3D chromatin organization changes in HSPCs and neutrophils

Having established HiCFoundation’s superior performance across multiple chromatin architecture analysis tasks, we next applied the model to analyze a challenging collection of Hi-C data. Hematopoietic stem and progenitor cells (HSPCs) give rise to all blood and immune cell lineages [45]. Lineage determination involves an interplay between epigenetic modifications, key transcription factors, and alteration of 3D genome architecture. Nuclear lamin B1 (LMNB1), a structural protein essential for maintaining the integrity and morphology of the nuclear envelope [46], alters 3D chromatin organization in HSPCs and causes Pelger-Hüet nuclear anomaly, a distinct abnormality in neutrophils resulting in mono- or bi-lobed nuclear morphology. However, global Hi-C analysis in these cells has been challenging due to the limited number of primary HSPCs, resulting in Hi-C contact maps with limited resolution [47]. Notably, the detection of chromatin loops, which are the most dynamic 3D genome feature and are closely associated with gene regulation, has been very challenging. To address these challenges, we used HiCFoundation to characterize dynamic changes in chromatin loop organization during HSPC differentiation into neutrophils. We first applied HiCFoundation-Reso to enhance the Hi-C matrices from control and LB1^*LO*^ human CD34+ HSPCs and neutrophils, where LB1^*LO*^ represents *LMNB1* knockdown resulting in low level (∼25%) of lamin B1 expression [47]. The enhancement substantially improved overall coverage and reduced noise while preserving relative coverage differences across samples (**Extended Data S7a**). The enhanced data was further validated by comparing different TAD insulation score patterns across four samples, where we observed similar patterns between raw and enhanced data (**Extended Data S7e**). Enhancing the Hi-C data increased differences in strength between the boundary and off-boundary regions, which helped TAD identification and differential analysis. This analysis revealed that TAD boundary strength is lower in neutrophils compared to HSPC and also upon lamin B1 loss in both cell types.

The enhanced, high-resolution Hi-C matrix resulted in the identification of chromatin loops that were previously undetected from the low-coverage data (**Fig. 4a**). We were able to detect as many as *>*7-fold more short-range chromatin loops across all samples. Notably, neutrophils had overall fewer chromatin loops across the genome compared to HSPCs, a difference that was also observed in the unenhanced data but with low confidence given the unreliable loop calling (**Fig. 4b**). The distribution of loop lengths was similar across all samples, suggesting that the relative distribution of short-range, mid-range, and long-range chromatin interactions remains unchanged (**Extended Data S7b**,**c**). The enhanced dataset confirms that lamin B1 loss results in both loop loss and loop gain; however, interestingly, the total number of loops in LB1^*LO*^ HSPCs or neutrophils was not substantially altered (**Fig 4b, Extended Data S7d**). We next compared loop strength as measured by peak-to-lower-left ratios (P2LL, see **Methods**) [48, 49]. This analysis showed a reduction in loop strength in neutrophils relative to HSPCs, and as a result of lamin B1 loss in both HSPCs and neutrophils (all sign tests yield *p <* 0.001, **Fig. 4c**). The decrease in strength is consistent across all loops as well as in comparisons of conserved loops. Aggregate peak analysis across samples revealed patterned changes in loop architecture (**Fig. 4d**). In particular, we observed that LB1^*LO*^ neutrophils exhibit the lowest loop strength, primarily due to stronger signals in the lower-left region. Furthermore, the loop peak signal in control neutrophils is significantly weaker than that in control HSPCs on shared loops. These enhanced data thus reveal a global loss of short-range chromatin loops in neutrophils, consistent with the finding that halting loop extrusion is essential for neutrophil differentiation [50]. Moreover, our findings show that lamin B1 loss results in global weakening of loop strength in both HSPCs and neutrophils.

**Figure 4:**
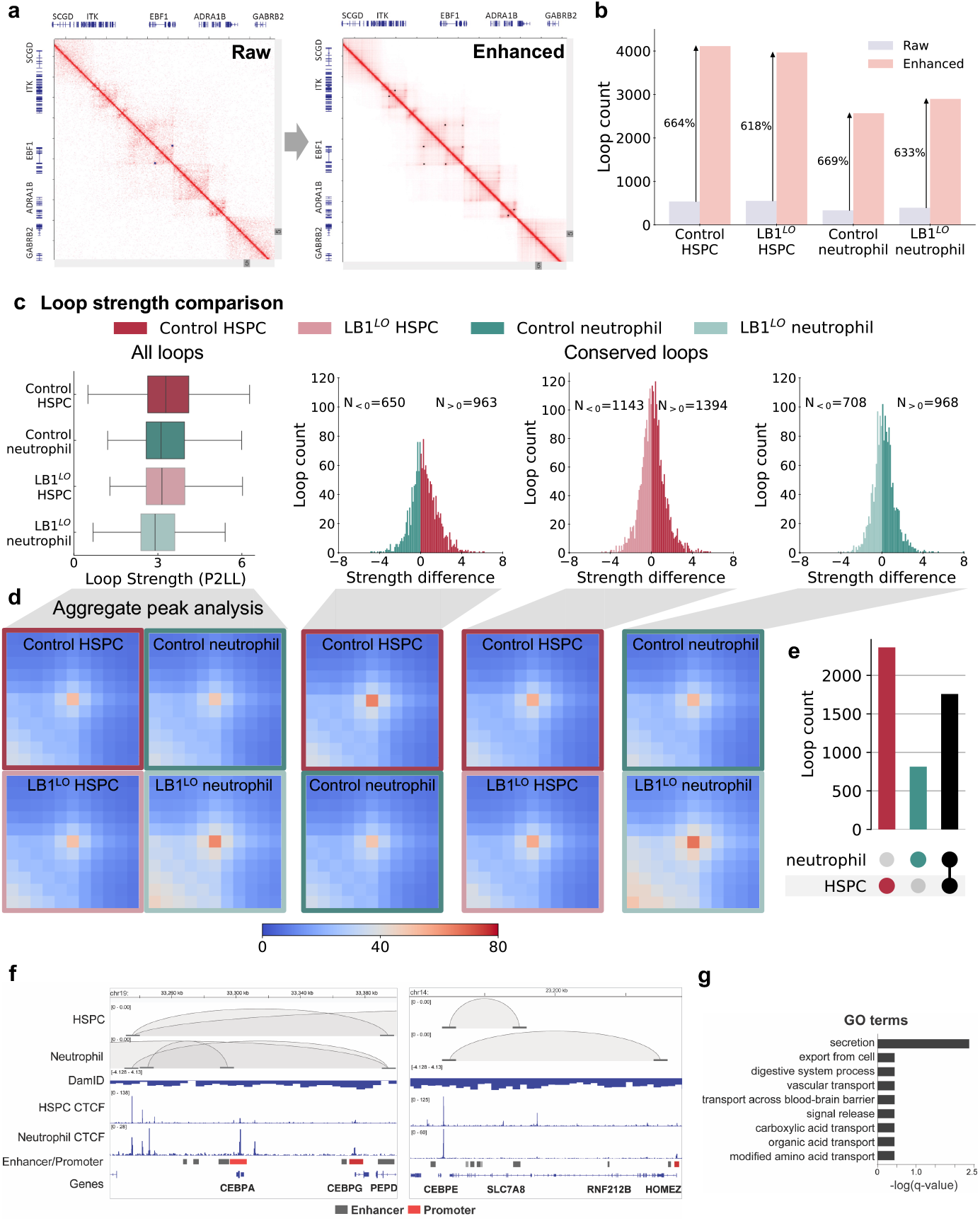
3D chromatin architecture changes revealed by HiCFoundation in HSPCs and neutrophils. The comparisons are made using four samples: control HSPC, LB1^*LO*^ HSPC, control neutrophil, and LB1^*LO*^ neutrophil. **a**. Representative Hi-C contact matrices on low coverage raw data and post-enhancement (blue dots indicate annotated loops). **b**. Loop counts in all samples on raw and enhanced data. **c**. Loop strength (peak to lower-left ratios, P2LL) comparison across samples via enhanced Hi-C. The left panel compares all loops across four samples. The right three panels compare the loop strength on conserved loops, defined as loops called in all four samples. **d**. Aggregate peak analysis of chromatin loops across samples. The left panel compares loop strength of all loops across four samples. The right panel presents the loop strength of conserved loops. The three comparisons are the same as panel c. **e**. Upset plot of gene-annotated loops from control HSPC and neutrophils. A gene is assigned to a loop if it is entirely within one of the associated 10 kb loop anchors. **f**. Chromatin loops observed at the *CEBPE* and *CEBPA* loci aligned to CTCF ChIP-seq from neutrophil (GSE101279), CD34+ HSPC, and GeneHancer track [43]. **g**. Gene Ontology analysis using Metascape [44] on genes associated with loops having higher strength in neutrophils vs. HSPC.

Although neutrophils predominantly lose chromatin loops that were present in HSPCs, we found that 806 new loops were gained in neutrophils compared to HSPCs (**Fig. 4e**). We hypothesized that these neutrophil-specific loops correspond to genes essential for neutrophil cell identity and function. To this end, we first examined looping at genes essential for neutrophil differentiation, including the C/EBP-family TFs *CEBPA* and *CEBPE* (**Fig. 4f**). At both loci, we observed reconfiguration of enhancer-promoter loops to alternative distal or proximal enhancers, corresponding with increased gene expression. We next performed an unbiased Gene Ontology analysis on loops that gained loop strength in neutrophils (log2(P2LL_HSPC vs. neutrophil_ *>*0.5)), which revealed a significant enrichment of genes involved in secretion, including *AQP5, CHRM1, LTBP4, ABCC4* and *TSPAN18* (**Fig. 4g**). Taken together, our enhanced dataset shows that although neutrophil differentiation is accompanied by loss of loops genome-wide, likely reflecting global chromatin compaction, looping around genes essential for neutrophil function is maintained and reconfigured to permit their continued expression. Moreover, lamin B1 may contribute to neutrophil differentiation by stabilizing loop configurations, with lamin B1 depletion resulting in weakening of loop strength.

### HiCFoundation profiles epigenomic assays

One of the main incentives for interpreting 3D genome architecture is to understand the regulatory implications of DNA 3D organization. Integrative analyses combining Hi-C with assays measuring local chromatin accessibility, TF binding, and histone modification profiles have revealed how 3D genome architecture influences gene activity [1]. For example, loop interactions identified by Hi-C are significantly enriched with cis-regulatory elements, particularly active promoters, enhancers, and CTCF binding sites [51, 52]. Leveraging this relationship between 1D and 3D organization, a recent deep learning model, Epiphany [53], has demonstrated the feasibility of predicting Hi-C contact maps from a collection of epigenomic measurements, including DNase I hypersensitive sites and CTCF, H3K27ac, H3K27me3, and H3K4me3 ChIP-seq. However, because the resolution of Hi-C data is often much lower than that of other epigenomic signals, predicting in the reverse direction—that is, predicting epigenomic signals using Hi-C data—remains challenging.

We hypothesized that HiCFoundation could tackle this resolution disparity problem by aggregating the chromatin structure patterns from a large quantity of Hi-C data, even though these data are at a relatively low resolution. We therefore created HiCFoundation-epi to generate a collection of 1D epigenomic measurements by using Hi-C data as input, thereby allowing us to study the effects of chromatin contacts on gene regulation. To this end, we fine-tuned HiCFoundation-epi using a submatrix of 128 kb by 4 Mb at 1 kb resolution, to produce six epigenomic signals for the corresponding 128 kb region, including two measurements of chromatin accessibility (DNase-seq and ATAC-seq), CTCF ChIP-seq, and three histone ChIP-seq profiles (H3K4me3, H3K27ac, and H3K27me3). The resulting model can thus generate genome-wide epigenomic profiles of these six assays (**Fig. 5a**, see **Methods**).

**Figure 5:**
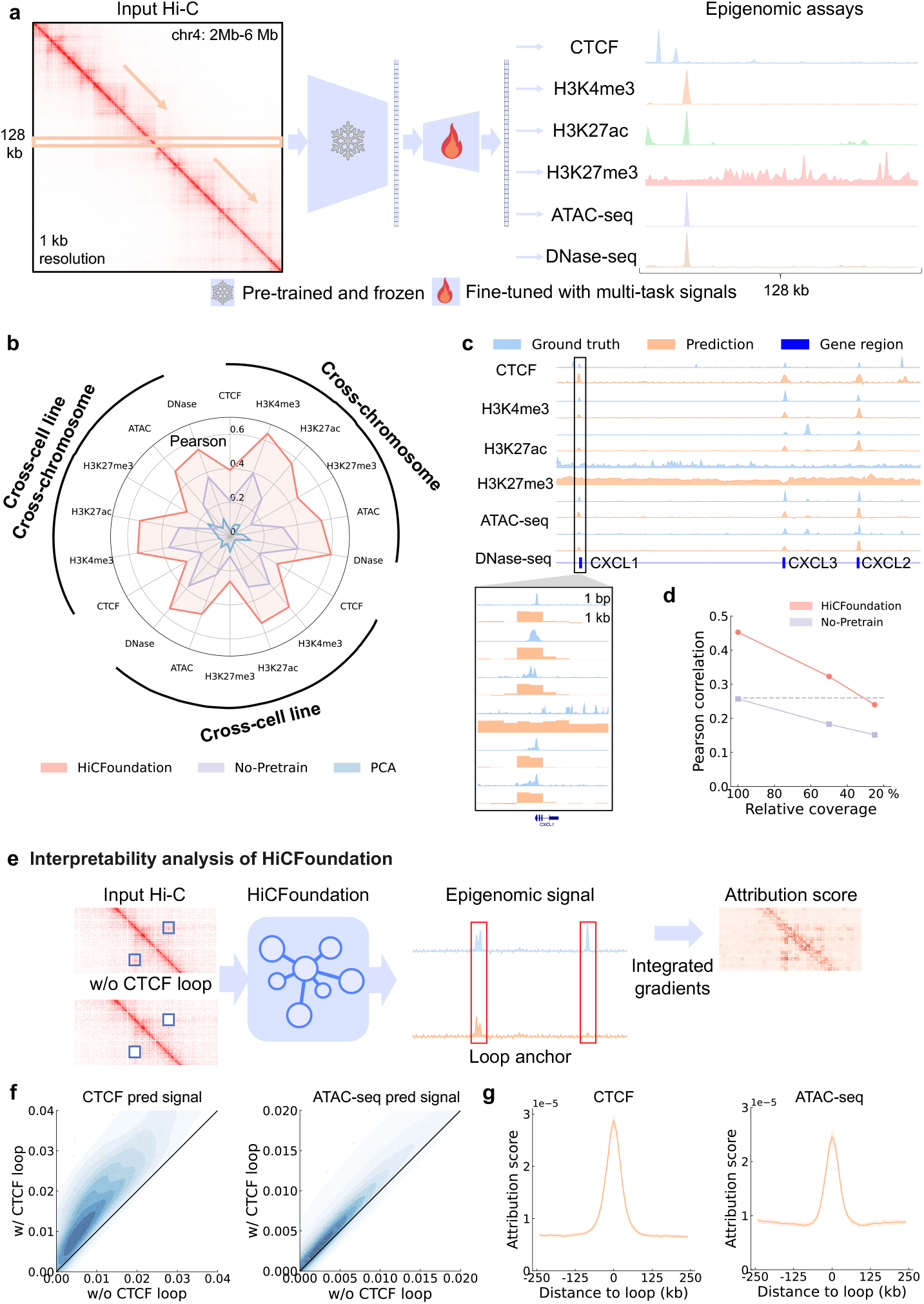
HiCFoundation for profiling epigenomic assays. **a**. HiCFoundation takes as input a raw 128kb ×4Mb Hi-C matrix and outputs the corresponding six 128kb epigenomic assays at 1kb resolution. The decoder is fine-tuned via multi-task training. **b**. Benchmark of HiCFoundation for epigenomic assays profiling. Each axis corresponds to the pearson correlation of different epigenomic assays across three different test sets: cross-chromosome, cross-cell line, and cross-chromosome and cross-cell line. **c**. Visualization of six epigenomic signal predictions on the K562 cell line in the CXCL1–CXCL3 gene region. A zoomed in view compared the predictions at 1 kb resolution and the experimental measurement at 1 bp resolution. **d**. Performance of HiCFoundation across varying Hi-C input coverage levels. **e**. Interpreablility analysis of HiCFoundation. That includes two settings to study the impact of CTCF loops on epigenomic assay profiling. In the first setting, for each loop, we compare two versions of the predicted epigenomic signals: one derived from the original Hi-C matrix and the other from the Hi-C matrix after masking out the CTCF loop regions. In the second setting, we analyze the attribution score of HiCFoundation for the input Hi-C using integrated gradients. This interpretation algorithm assigns attribution scores to the Hi-C input, indicating the relative importance of each contact within the Hi-C matrix. **f**. CTCF ChIP-seq and ATAC-seq predicted signal with and without CTCF loops. **g**. Attribution score distribution for CTCF ChIP-seq and ATAC-seq predictions relative to the distance from the loop. Values are averaged over 392 loops, and error bars correspond to standard deviation. Dataset details are available in **Supplementary Table 11**, and the benchmark of different methods is in **Supplementary Table 12**.

HiCFoundation-epi was fine-tuned on data from two human cell lines, GM12878 and H1ESC, and then evaluated in three distinct settings: 1) cross-chromosome testing on GM12878 and H1ESC; 2) cross-cell line testing on K562; and 3) cross-cell line and cross-chromosome testing on K562. To the best of our knowledge, there is no existing method that can systematically predict epigenomic profiles from Hi-C data. Therefore, we compared HiCFoundation-epi to two baselines: No-pretrain and HiCPCA [54], which uses principal component analysis (PCA) to reduce the 2D Hi-C matrix to a 1D track. HiCFoundation-epi consistently yields superior performance against these baselines in all three settings (**Fig. 5b, Supplementary Table 12**). For example, HiCFoundation-epi achieves Pearson correlations of 0.294, 0.549, 0.537, 0.292, 0.500, 0.544 for CTCF, H3K4me3, H3K27ac, H3K27me3, ATAC-seq, and DNase-seq, respectively, representing an average improvement of 0.192 (min=0.103, max=0.255) relative to No-pretrain and 0.365 (min=0.141, max=0.485) relative to HiCPCA. We further evaluated HiCFoundation’s performance under varying coverage levels by downsampling. HiCFoundation achieved comparable performance to the No-pretrain model with only around 25% of the original coverage (**Fig. 5d**). This demonstrates HiCFoundation’s robustness across different coverage levels and highlights its predictive power even with shallow read depth, reducing both experimental complexity and financial cost.

Visual examination of chromosome 4 in K562 cells (**Fig. 5c**) suggests that HiCFoundation-epi yields qualitatively accurate predictions, even in the most challenging cross-cell line and cross-chromosome setting. Specifically, we observed strong consistency between the predicted and measured epigenomic signals at the CXCL1–CXCL3 locus. These genes, which encode chemokines from the CXC family, serve as critical regulators of immune and inflammatory responses [55]. Notably, data from the BBcancer database [56] reveal significant expression differences in these genes between cancer patients and normal controls. Consequently, accurate profiling of the associated epigenomic signals is essential for enabling robust downstream analyses, which also highlights the great potential of HiCFoundation.

Given the strong performance of HiCoundation-epi, we hypothesized that the model should be able to reveal the relationship between genome architecture changes and corresponding epigenomic signal changes. To validate this hypothesis, we exploit the known association between CTCF ChIP-seq peaks and Hi-C loops, which arises due to cohesin-mediated chromatin loop formation [51, 52]. In particular, we first used HiCFoundation-loop to detect loops in test set chromosomes from the K562 cell line, and we restricted our analysis only to the 392 loops that contain convergent CTCF motifs identified using FIMO [57]. Next, for each loop, we compared the two versions of the predicted CTCF ChIP-seq signals at the loop anchor. One is predicted using the original contact matrix; the other is predicted using the contact matrix that zeros out the 30 ×30 kb region around the loop (**Fig. 5e**). This comparison tests whether HiCFoundation-epi can correctly capture the change in CTCF ChIP-seq signals due to the presence of CTCF loops.

We found that in 97.8% (383 out of 392) of cases, the ablation of the loop leads HiCFoundation-epi to predict decreased CTCF ChIP-seq signal, with an average decrease of 41.8% (**Fig. 5f**). The same relationship is observed between the predicted ATAC-seq signal and CTCF loops using a similar analysis. To further assess the generated profiles, we used Integrated Gradients [12] to assign an attribution score to each contact in the input locus Hi-C (**Fig. 5e**). We observed that the attribution score of a contact is strongly correlated with its distance to the loop, indicating that HiCFoundation-epi used the correct contact information to predict epigenomic signals (**Fig. 5g**). These two interpretability analyses of HiCFoundation-epi validate its utility for investigating relationships between genome architecture and epigenomic assay profiling, paving the path for integrative studies of gene regulation.

### HiCFoundation generalizes to single-cell Hi-C analysis

Hi-C captures DNA–DNA contacts from cell populations, producing a population average 3D genome structure that may not reflect 3D genome architecture in individual cells. Accordingly, scHi-C was developed to measure chromatin architecture within a single nucleus [3]. However, its coverage is orders of magnitude lower than bulk Hi-C, resulting in a low signal-to-noise ratio and posing challenges for downstream analysis [61]. Thus, resolution enhancement is particularly valuable for scHi-C analysis. To address this challenge, several scHi-C resolution enhancement methods have been developed [62, 63, 64]. However, these methods are unsupervised methods that rely on certain assumptions on the contact distribution, such as that nearby loci exhibit similar contacts, which might lead to less accurate performance on cells with low coverage. Furthermore, many of these methods work by borrowing information from neighboring cells (i.e., cells with similar patterns of scHi-C data), which potentially reduces the true cell-to-cell variability in the data.

We hypothesize that HiCFoundation can leverage patterns from bulk Hi-C data to enhance resolution for scHi-C by fine-tuning on scHi-C data. To this end, we fine-tuned HiCFoundation-sc for scHi-C resolution enhancement using scHi-C data from 6060 single cells from developing mouse embryos and brains, generated using the HiRES scHi-C/scRNA-seq co-assay (HiRES dataset) [58]. The HiCFoundation-sc model takes as input a full chromosome of scHi-C data at 1 Mb resolution and then outputs a corresponding enhanced matrix (**Fig. 6a**). The scHi-C-specific decoder is optimized using pairs of raw scHi-C data and 4 times-downsampled scHi-C data. To cope with the extreme sparsity of scHi-C data, we propose a novel rank-based MSE loss that effectively reduces the sensitivity of the loss gradients to large values (see **Methods** and **Extended Data S8a**).

**Figure 6:**
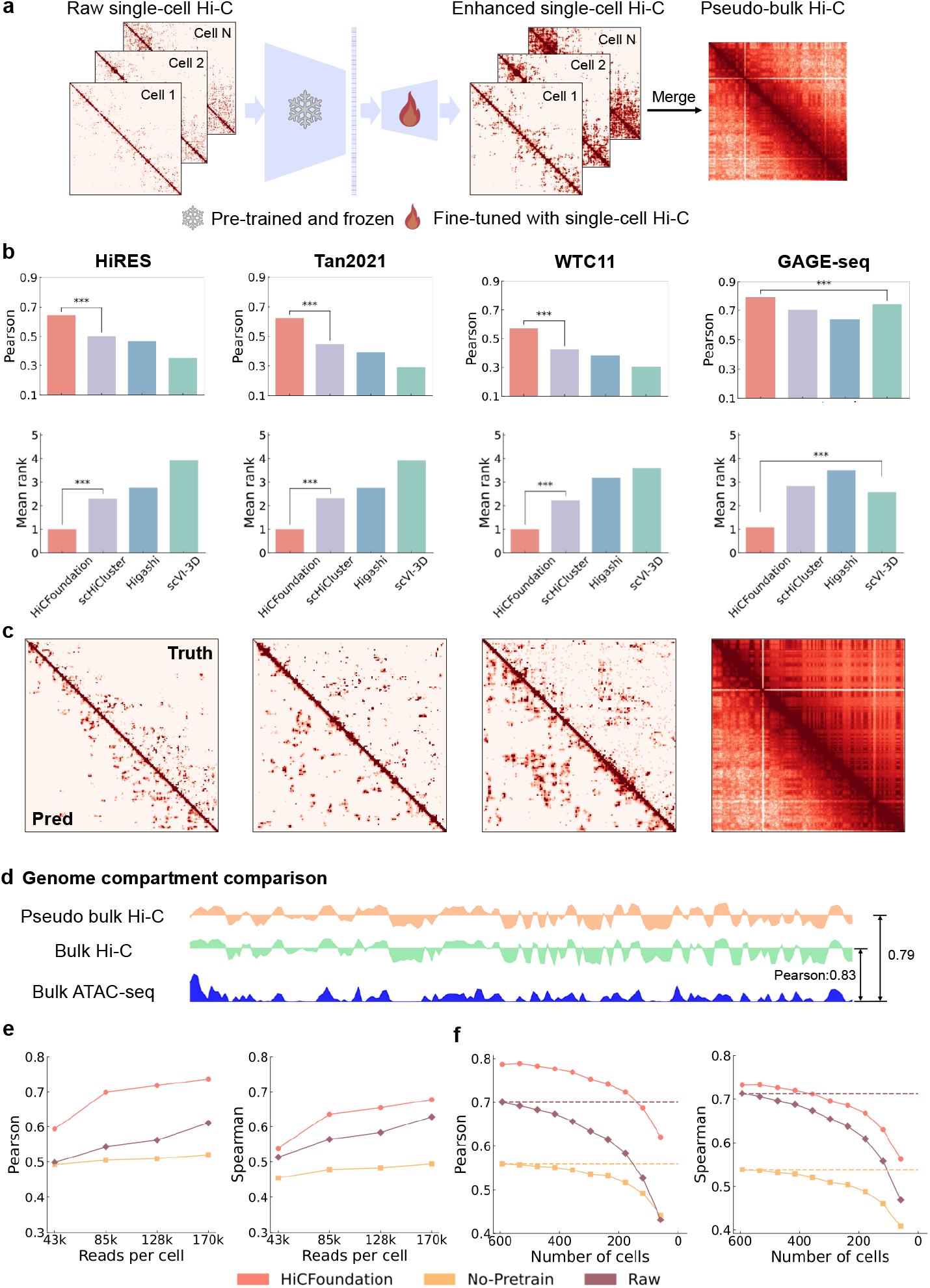
HiCFoundation for single-cell Hi-C analysis. **a**. Inference pipeline of HiCFoundation-sc on single-cell Hi-C resolution enhancement. HiCFoundation-sc can takes chromosome scHi-C matrix as input and output the enhanced scHi-C matrix. This can be further merged into pseduo-bulk Hi-C to compare against bulk Hi-C. **b**. Benchmark on four different datasets of four settings: 1) cross-cell and cross chromosome setting on developing mouse embryos cells HiRES dataset [58]. 2) cross-chromosome and cross-dataset testing on mouse forebrain cortex cells [59]. 3) cross-species testing on data from the human cell line WTC11 [14]. 4) pseudo-bulk and bulk Hi-C benchmark on human cell line K562 [60]. The other evaluation metrics are included in **Extended Data S8b-e c**. Visual comparisons between enhanced matrices (lower left) generated by HiCFoundation-sc and the ground truth (GT) (upper right). The left three panels display three randomly selected single-cell enhancements and their corresponding ground truth from the HiRES, Tan2021, and WTC11 datasets. The right panel shows pooled enhanced scHi-C (pseudo-bulk) compared to bulk Hi-C from the GAGE-seq dataset. **d**. Genome compartment detection example. We compared the correlation between the PCA of pseudo-bulk Hi-C by HiCFoundation-sc and bulk Hi-C with the corresponding bulk ATAC-seq measurements. **e**. Performance comparison of various approaches across different reads per cell on GAGE-seq dataset. **f**. Performance of HiCFoundation-sc across varying single-cell dataset sizes (number of cells) on GAGE-seq dataset. The HiCRep evaluations of panel e and f are included in **Extended Data S9e,f** The dataset information is available at **Supplementary Table 13** and the benchmark performance is available at **Supplementary Table 14**. “***” in the panels indicates significance based on the sign test with (*p <* 10^−3^).

We evaluated HiCFoundation-sc in three settings: 1) cross-chromosome testing using held-out chromo-somes from the HiRES dataset; 2) cross-chromosome and cross-dataset testing on mouse forebrain cortex cells [59]; and 3) cross-species testing on data from the human cell line WTC11 [14] (**Supplementary Table 13**). In each setting, performance is evaluated by comparing the enhanced scHi-C data to the original scHi-C data. Across all three settings, HiCFoundation consistently delivers superior performance compared to three state-of-the-art methods, Higashi [62], scHiCluster [63], and scVI-3D [64] across various metrics (**Fig. 6b Extended Data S8b-d, Supplementary Table 14**). Moreover, HiCFoundation yields the largest improvement in the most challenging cross-species setting, where it attains 35.9% improvement on Pearson correlation, 20.0% improvement on SSIM, and 30.2% improvement in PSNR over the second best method, scHiCluster (all sign tests, *p <* 0.001). For this setting, HiCFoundation is directly applied without training on WTC11 data, whereas all other methods are retrained on the WTC11 dataset, further validating the generalizability and power of HiCFoundation.

We further validated the generalizability of HiCFoundation-sc using human K562 cell line data generated in the GAGE-seq co-assay [60]. Rather than evaluating on downsampled data, this evaluation compares pseudo-bulk from enhanced scHi-C against bulk Hi-C data from the same cell line (**Fig. 6b**). HiCFoundationsc is directly applied without training, whereas all other methods are trained directly on the GAGE-seq dataset (see **Methods**). Due to coverage differences between pseudo-bulk and bulk Hi-C data, SSIM and PSNR were excluded as evaluation metrics. We found that HiCFoundation-sc achieves the best performance across metrics, with a Mean rank of 1.167, which is substantially better than competing methods (right-most panels of **Fig. 6b**). A visual comparison between HiCFoundation-sc and ground truth across four datasets further highlights its strong enhancement capability (**Fig. 6c**). We further applied HiCPCA [54] to the merged enhanced scHi-C data (pseudo-bulk Hi-C) and bulk Hi-C data on GAGE-seq dataset. We observed that the compartments detected by the pseudo-bulk Hi-C show strong consistency with the compartments revealed by the corresponding ATAC-seq measurements (**Fig. 6d**), comparable to the compartments observed from bulk Hi-C.

Following the same setting as before, we also compared the performances of HiCFoundation relative to No-pretrain for scHi-C resolution enhancement (**Extended Data S9**). The relatively poor performance of No-pretrain reflects the di”culty of applying supervised models across datasets with varying coverages. In contrast, HiCFoundation-sc consistently delivers strong results with a large margin over other approaches, underscoring the strong generalization of HiCFoundation’s embeddings across diverse assays, even at single-cell resolution.

Lastly, we evaluated the robustness of HiCFoundation-sc across varying coverage levels and numbers of individual cells on the GAGE-seq dataset. First, we curated a fixed-coverage scHi-C dataset with uniform reads per cell by discarding 50% of the cells with reads below the 50th percentile and downsampling the remaining cells to equal read counts. Next, we compared the performance of pseudo-bulk data derived from raw data, No-pretrain, and HiCFoundation-sc under varying coverage levels by downsampling the fixed-coverage dataset (**Fig. 6e** and **Extended Data S9e**). Compared to raw and No-pretrain, we observed that HiCFoundation’s performance is stable with more than 85k reads per cell, which suggests the robustness of HiCFoundation. Next, we investigated the performance of HiCFoundation with different numbers of cells in the dataset (**Fig. 6f** and **Extended Data S9f**). Notably, HiCFoundation requires only 150 cells to generate a pseudo-bulk dataset with performance comparable to that achieved using around 600 cells from raw scHi-C data. Together, these results validate the stability and effectiveness of HiCFoundation for scHi-C analysis, highlighting its potential in cases where achieving su”cient coverage or obtaining an adequate number of single cells is experimentally challenging.

## Discussion

HiCFoundation generates various types of embeddings that are broadly useful for a wide variety of Hi-C-related analysis tasks. Through extensive benchmarking, we show that HiCFoundation consistently delivers state-of-the-art performance across diverse tasks. Furthermore, fine-tuned models derived from HiCFoundation can be integrated in a unified framework for genome architecture and epigenomic functional analysis. For example, HiCFoundation-Reso and HiCFoundation-Loop can be combined to accurately detect chromatin loops from low-coverage Hi-C data, including human HSPCs and other rare primary cell types. Meanwhile, HiCFoundation-epi enables the profiling of chromatin accessibility, transcription factor binding, and histone modifications. Interpretability analysis of HiCFoundation-epi reveals its ability to effectively capture epigenomic signal at chromatin loop anchors. HiCFoundation thus provides a unified framework for Hi-C analysis, facilitating integrative, multi-species, multi-omics, and single-cell studies. With its streamlined and e”cient fine-tuning pipeline, this framework can be applied to a wide range of genomic and epigenomic analyses.

The most compelling advantage of HiCFoundation is its strong adaptability, which is facilitated by three factors. First, adapting HiCFoundation to new tasks is time e”cient. The pre-training of HiCFoundation from hundreds of Hi-C experiments is time-consuming, requiring approximately two weeks on a server equipped with 8 A100 GPUs. In contrast, fine-tuning HiCFoundation on a specific downstream tasks takes less than 10 hours due to HiCFoundation’s asymmetric encoder-decoder architecture, where the decoder is much smaller than the encoder. Only parameters of the lightweight decoder are updated using task-specific labels, while the encoder remains frozen during fine-tuning. Second, adapting HiCFoundation to new tasks is “label e”cient,” meaning that only a small amount of labeled data is required. This e”ciency allows the model to achieve substantial improvement over models trained from scratch, as demonstrated by our experiments. Third, HiCFoundation can produce various types of embeddings, including locus embeddings, patch embeddings, submatrix embeddings, and chromosome embeddings, making the model applicable to genomic and epigenomic analyses at different scales.

Hi-C datasets of human HSPCs have low coverage, limiting detection of chromatin loops and biological insights into gene regulation. By applying HiCFoundation to a dataset of human HSPCs and neutrophils, we improved detection of chromatin loops by over 7-fold. These data have uncovered a dramatic loss of chromatin loops and weakening of loop strength during neutrophil differentiation, while looping at genes essential for neutrophil function is preserved. Chromatin in neutrophils undergoes profound compaction as it is partitioned into 3–5 distinct nuclear lobes [65]. Loss of chromatin loops may be essential to achieve a high degree of physical compaction, consistent with the recent finding that halting loop extrusion is essential for neutrophil differentiation [50].

Although HiCFoundation already provides excellent performance across a variety of downstream tasks, some avenues for future research remain open. First, HiCFoundation currently does not incorporate sequence information, which might enhance its accuracy and utility for genome structure and functional analysis. Prior studies [66, 67, 68, 69, 70, 11] have shown the power of sequence-based approaches in predicting chromatin contacts and epigenomic assays. Integrating sequence information into HiCFoundation should further improve its performance. Second, although HiCFoundation excels at predicting epigenomic assays from Hi-C data, we hypothesize that bidirectional translation between Hi-C and epigenomic data, perhaps by integrating with a model like Epiphany, could allow the model to improve on both of these translation tasks.

## Acknowledgements

The authors thank Xinlei Chen, Gang Li, Ran Zhang, Justin Sanders, Bo Wen, Addie Woicik, Zucks Liu, Hanwen Xu, and Yu Zhang for discussions and suggestions for this paper. Several components of Fig. 1 are made using BioRender: https://biorender.com.

This work was partly supported by the National Institutes of Health (UM1 HG011531, R01 HG013321 and R21 EB036205). SW is supported by a Sony Faculty Research Award. XW is a UW Data Science Postdoctoral Fellow. SD and WSN are supported by NHLBI R01 HL169156. SD is supported by grants from NHLBI (R01 HL151651), NIDDK (RC2-DK127989), and Edward P. Evans Foundation. SD is a Scholar of the Leukemia and Lymphoma Society (1391-24).

## Author Contributions Statement

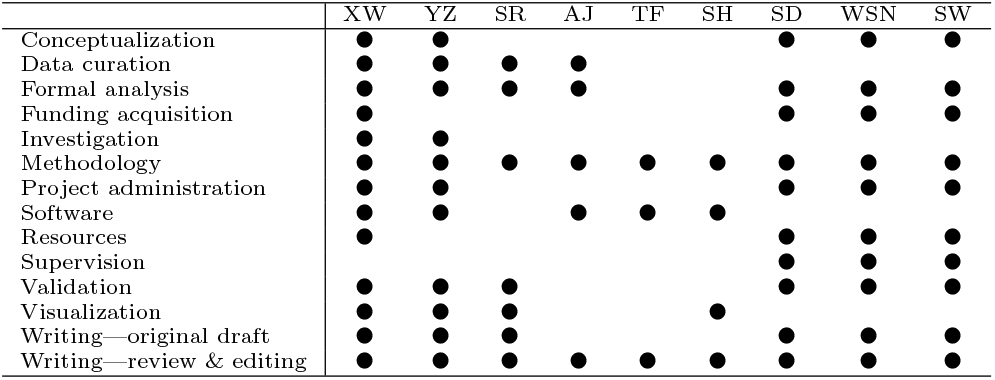

## Competing Interests Statement

Authors declare that they have no competing interests.

## Data Availability

The Hi-C and epigneomic assay data is downloaded from ENCODE (https://www.encodeproject.org/), 4D Nucleome (https://data.4dnucleome.org/) and DNA Zoo (https://www.dnazoo.org/). The accession IDs used in pre-taining and the downstream are listed in **Supplementary Table 1, 3, 5, 7, 10, 11**. The HSPC and neutrophil related data is available at https://www.ncbi.nlm.nih.gov/geo/query/acc.cgi?acc=GSE174533. For single-cell Hi-C datasets, the WTC-11 dataset is available with accessions 4DNESF829JOW and 4DNESJQ4RXY5 at the 4D Nucleome web portal (https://data.4dnucleome.org/); the Tan2021 dataset is available at https://www.ncbi.nlm.nih.gov/geo/query/acc.cgi?acc=GSE223917; the HiRES dataset is available at https://www.ncbi.nlm.nih.gov/geo/query/acc.cgi?acc=GSE119171; and the GAGE-seq dataset is available at https://www.ncbi.nlm.nih.gov/geo/query/acc.cgi?acc=GSE238001. The cell IDs used in our experiment are listed in **Supplementary Table 13**. The data processing code is available at https://github.com/Noble-Lab/HiCFoundation_paper.

## Code Availability

The HiCFoundation source code available https://github.com/Noble-Lab/HiCFoundation with an Apache license. The code, as well as pre-trained and fine-tuned models, are also available on Zenodo at https://doi.org/10.5281/zenodo.14436390. We also provide notebook tutorials on Github.

## Methods

### Pre-training data sets

We began by collecting a large set of Hi-C experiments, from which we created a pre-training set and a test set, as well as train and test sets for each of the three downstream tasks.

The full dataset consists of 678 Hi-C experiments generated by two National Institutes of Health consortia, ENCODE [13] and 4D Nucleome [14]. We used contact maps generated using the *in situ* Hi-C [71], dilution Hi-C [2], intact Hi-C, Micro-C [72], and DNase Hi-C [73] protocols. The data are derived from five different organisms: human, mouse, chicken, zebrafish and fruit fly. We only included in our dataset Hi-C experiments that contain at least 10 million non-diagonal mapped read pairs. We use the ENCODE terminology “biosample” to refer to the cell line or tissue type from which the sample is derived. Our dataset contains Hi-C data from 146 distinct biosamples. The Hi-C experiments vary substantially in the total number of read pairs, with a trend of increasing coverage over time. The full set of accession codes is provided in **Supplementary Table 1**.

We split this dataset into training, validation, and test sets. The splitting is done at the level of (human) biosamples, with 81 used for training, 20 for validation, and 18 for testing. This split corresponds to 368 Hi-C experiments in the training set, 36 in the validation set, and 117 in the test set. In addition, all non-human Hi-C data, consisting of 157 experiments, are used for testing. **Supplementary Table 1** shows which subset (training, validation, or test) each Hi-C experiment was assigned to. The training set is also referred to as the “pre-training set” in the literature of foundation models. The validation set is used during training to select the best-performing model, and the test set is used to evaluate model generalizability over unseen data.

We use a two-step procedure to prepare each Hi-C experiment for input to our model. First, we partition the collection of Hi-C matrices into a set of pairwise chromosome matrices. Each pairwise matrix comes from a single Hi-C experiment and contains data from two distinct chromosomes, excluding the Y chromosome. This design ensures that there is no information loss, as both intra-chromosomal and inter-chromosomal information is preserved within the pairwise chromosome Hi-C matrices. In total, we obtain 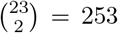 chromosome pairs per Hi-C experiment; thus, the training and validation sets yield a collection of 253× 404 = 102, 212 pairwise chromosome Hi-C matrices for pre-training. Second, each pairwise matrix is decomposed into 224 ×224 submatrices at 5 kb resolution. Submatrices that overlap chromosome boundaries are excluded, as are submatrices with *>*95% zero values. This procedure produces ∼116 million submatrices for training and 19 million submatrices for validation. In practice, during each training and validation epoch we randomly sample 1.16 million submatrices for training and 0.2 million submatrices for validation.

### Benchmarking data sets

For the Hi-C reproducibility task, we select a subset of the training set for training and a subset of the test set for testing. For training, we begin by selecting 109 experiments that have at least two biological replicates. Of these 109, we eliminate 80 experiments in which at least one of the two replicates has fewer than 10 million non-diagonal mapped read pairs. The remaining 29 experiments are drawn from 29 distinct human biosamples, which we randomly split into 24 for training and 5 for validation. In the ENCODE and 4DN repositories, each Hi-C experiment consists of at least two biological replicates. For each such experiment, we create one matrix from each biological replicate (BR). The training set thus consists of 24 BR pairs and 1104 non-replicate (NR) pairs, and the validation set consists of 5 BR pairs and 40 NR pairs. For the test set, we apply a similar procedure, yielding 15 BR pairs from human and 20 BR pairs from mouse, as well as 420 human NR pairs and 760 mouse NR pairs. Note that both BR and NR pairs are derived from single replicates, not pooled replicates as in the pre-training stage.

Similarly, for the Hi-C loop detection task we construct our dataset by selecting subsets from the training and test sets. We exclude experiments 1) with one biological replicate, 2) if either of the two biological replicates had fewer than 100 million non-diagonal mapped read pairs, or 3) if HiCCUPs called fewer than 1000 loops in either of the two replicates. This process results in 15 experiments sourced from 15 biosamples, including 6 from human and 9 from mouse. From these, we select 4 experiments from human biosamples that are in the training set of the pre-training stage to serve as the training and validation set for the loop detection task. In this set, chromosomes 4, 5, 11, and 14 are designated as the validation set, while all other chromosomes are used for training. The testing set includes chromosomes 4, 5, 11, and 14 from the remaining 2 experiments from human, plus all chromosomes from the 9 experiments sourced from mouse.

For resolution enhancement, we construct our dataset using subsets from the training and testing sets. To ensure that the down-sampled Hi-C data has reasonable read counts, we first filter out experiments with fewer than 500 million reads. This leaves us with 124 experiments for training and validation, and 27 experiments for testing. Among the training and validation experiments, chromosomes 4, 5, 11, and 14 are designated as the validation set, following the procedure used in HiCARN [33] and DeepHiC [74] to enable a fair comparison. All other chromosomes are used for training. Our testing set includes 16 experiments sourced from humans and 11 experiments sourced from mouse. To test the generalizability of our model, we benchmark it on chromosomes 4, 5, 11, and 14 from human, and on all chromosomes from mouse, which were never seen during training.

For the task of predicting 1D epigenomic signals from Hi-C, we construct our dataset using ENCODE and 4DN data, following the approach adopted by Epiphany [53]. Specifically, we gather seven types of data: Hi-C, ATAC-seq, and DNase-seq, as well as ChIP-seq for CTCF and three types of histone modifications (H3K4me3, H3K27ac, H3K27me3). All data types are collected from the GM12878, K562, and H1ESC cell lines. Accession codes can be found in **Supplementary Table 11**. To prevent any potential data leakage, all three cell lines used here are excluded from the pre-training stage training set. For training and validation of the 1D epigenomic predictor, we use Hi-C and the corresponding epigenomic signal data from GM12878 and H1ESC, excluding chromosomes 4, 5, 11, 14, and X. After training, we evaluate our model across three different testing settings: 1) cross-chromosome setting, testing on chromosomes 4, 5, 11, and 14 of GM12878 and H1ESC; 2) cross-cell line setting, testing on all chromosomes, except 4, 5, 11, 14, and X, of K562; and cross-cell line and cross-chromosome setting, testing on chromosomes 4, 5, 11, and 14 of K562.

For single-cell Hi-C analysis, we train, validate and test our model using four publicly available scHi-C datasets (see **Supplementary Table 13** for more information). For these datasets, we keep cells with more than 100,000 intra-chromosomal contact reads. After this filtering process, the WTC11 dataset [75] contains 185 cells, the Tan2021 dataset [59] contains 1,943 cells, the HiRES dataset [58] contains 7,576 cells, and the GAGE-seq dataset [60] contains 593 cells from the K562 cell line. Given the quality and size of each dataset, we selected the HiRES dataset for model training and validation. We train the model on all train chromosomes using 80% of the cells, validate the model on chromosomes 4, 5, 11, and 14 on the training cells, and test the model on chromosomes 2, 6, 10, and 12 using the testing cells. For the Tan2021 dataset, 1,943 mouse cells with chromosome 2, 6, 10 and 12 are used for testing in the cross-dataset, cross-chromosome setting. Furthermore, we used the test chromosomes 4, 5, 11 and 14 on human cells, similar to the other downstream tasks. For the WTC11 dataset, 185 human cells along with chromosome 4, 5, 11, and 14 are benchmarked for the cross-species experiment. For the GAGE-seq dataset, 593 K562 cells are used for comparison with bulk HiC, focusing on chromosomes 4, 5, 11 and 14. Detailed information about all four datasets is available in **Supplementary Table 13**.

### HiCFoundation pre-trained model architecture and pre-training details

The HiCFoundation model training can be divided into two stages: the pre-training stage, which uses self-supervised learning to train from a large collection of unlabeled data, and the fine-tuning stage, which adapts the pre-trained HiCFoundation model for different downstream tasks using task-specific labelled data.

The self-supervised pre-training involves randomly masking some parts of a given submatrix and then training the model to fill in the masked regions. In particular, for every training epoch, we randomly sample 1.16 million 224 ×224 submatrices. Hence, each submatrix spans 5 kb ×224 = 1.12 Mb. Each of these submatrices is divided into a collection of 196 non-overlapping 16 ×16 patches. A random selection of 75% of these patches is masked, leaving 25% unmasked patches as input to HiCFoundation. Because Hi-C data is symmetric along the diagonal, we apply symmetric masking for any submatrix that includes any diagonal regions. For the output, the complete set of 196 patches from the input submatrix is used as the reconstruction target.

Similar to other approaches optimized for reconstruction, we used an encoder-decoder architecture. In particular, we adopt the widely used masked auto-encoder architecture (MAE) [76, 77] as the backbone of HiCFoundation due to its strong performance on images [16]. As shown in **Fig. 1c**, the MAE first splits an image into patches and then randomly masks a large fraction of these patches. The encoder converts each unmasked patch to an embedding vector, and the masked patches are mapped to a shared, learnable embedding vector. The MAE adds a sinusoidal positional embedding to the embedding of each masked and unmasked patch. The resulting combined embeddings are then used as the input for the decoder to reconstruct the entire image, including the masked patches. After pre-training the MAE, the encoder is used to embed new Hi-C submatrices for various downstream tasks, while the decoder is replaced by a task-specific architecture. Accordingly, the MAE architecture often uses a large encoder with many parameters and a simple decoder to maximize the effectiveness of the encoder. Moreover, previous work has empirically found that masking a very large proportion of patches leads to better results, since this approach makes the pre-training task more challenging [16]. Here we masked 75% of the patches and optimized the model to reconstruct the masked patches. The model uses an asymmetric encoder-decoder architecture design, where we adopted a larger encoder (304M parameters) and a smaller decoder (26M parameters) to maximize learning capacity during pre-training while minimizing fine-tuning costs.

Along with the patch embeddings from the Hi-C submatrices, our model also takes two additional embeddings as input. Because the Vision Transformer architecture typically includes in its input a class token embedding [78] to represent the entire submatrix’s embedding, we also append an auxiliary dummy embedding to the encoder input. This class token embedding can be used for fine-tuning downstream tasks. Additionally, we added as another input embedding a sinusoidally encoded count of the total number of mapped reads pairs associated with the corresponding Hi-C experiment

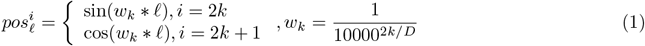

where 𝓁= log_10_(reads + 1), *D* is the feature dimension, and 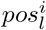 is *i*th element of vector *pos*_*l*_ [79, 78].

For the encoder, we employ the Vision-Transformer Large (ViT-L) architecture [78]. In this step, we define the 2D positional embedding 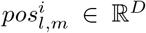 for the patch at row *l* and column *m* analogously to Equation 1:

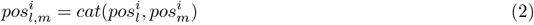

where 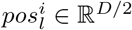 and 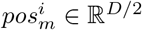 are the 1D positional embeddings of *l*th and *m*th patch, respectively.

We couple this encoder with a light-weight decoder consisting of eight transformer blocks with a latent dimension of 512. Note that the decoder is discarded after the pre-training stage, and only the pre-trained encoder is used in the subsequent fine-tuning procedures.

Many self-supervised learning frameworks for image reconstruction optimize an MSE loss between the reconstructed and original images in the pixel space [80, 16]. However, we found that the MSE loss applied to Hi-C data yields trivial solutions because of the high degree of sparsity in the data. Instead, the HiC-Foundation pre-training procedure optimizes a two-component loss function, with one component akin to a cross-entropy loss and a second component modeled after the structural similarity index measure (SSIM) [19].

Our first loss term is a contrastive loss function, called InfoNCE [81], applied at the level of patches:

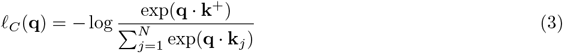

where *N* is the number of patches output by the model, and *q* is a given patch’s embedding, *k*^+^ is the embedding of the corresponding patch at the same row and the same column in the ground truth matrix. Here all embeddings are already L2 normalized. This loss function captures the intuition that *q* should be similar to its corresponding patch in the ground truth *k*^+^ and dissimilar to all other patches in the ground truth. The InfoNCE loss can be conceptualized as a variant of cross-entropy loss. The similarity is subsequently normalized by the softmax function to yield a pseudo-probability.

Whereas Equation 3 primarily focuses on local similarities, the second term of our loss function aims to enhance the global structural similarity between the reconstructed Hi-C submatrix and the ground truth Hi-C submatrix. This term is modeled on the SSIM:

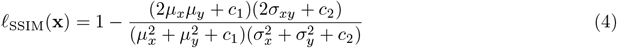

where *x* is the reconstructed Hi-C submatrix; *y* is the ground truth Hi-C submatrix; *µ*_*x*_ and *σ*_*x*_ are the pixel sample mean and variance of *x*; *µ*_*y*_ and *σ*_*y*_ are the pixel sample mean and variance of *y*; *σ*_*xy*_ is the covariance of *x* and *y*; and *c*_1_ and *c*_2_ are two constants that stabilize the division.

Combining the two loss terms, the HiCFoundation model is optimized by

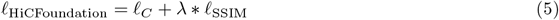

where *λ* is a hyperparameter that balances the two loss terms. In this work, we fix *λ* = 1.

The overall network is optimized using the AdamW [82] optimizer with an initial learning rate of 2.4 × 10^−3^, weight decay of 0.05, a batch size of 4096 and momentum parameters of *β*1 = 0.9 and *β*2 = 0.95. Due to limitations in available GPU resources, gradient accumulation [83] was employed to achieve a large batch size. The learning rate follows a decay schedule during the training process based on a cosine annealing schedule [84]. The pre-training phase consists of 100 epochs, with 5 epochs designated as warm-up epochs [85]. During the warm-up epochs the learning rate is increased in a linear fashion at each iteration such that the desired learning rate (2.4 × 10^*−*3^) is achieved after 5 epochs. This learning rate annealing approach is commonly used in training deep learning models to prevent instability or divergence in the early stages of training. After each epoch, we evaluated the model using the validation set, and we selected the model with the minimum validation loss as our final model for downstream tasks. The model was trained on a server equipped with 8 A100 GPUs, each with 80GB of memory, and the pre-training process required approximately two weeks.

### Fine-tuning HiCFoundation for the reproducibility task

For the reproducibility task, we use the pre-training encoder-decoder architecture. Specifically, we transfer the encoder weights from the pre-trained final model, while initializing the decoder weights via xavier uniform initialization [86]. During the fine-tuning stage, we keep the encoder weights frozen and only fine-tune the decoder weights.

Training the model to measure Hi-C reproducibility requires creating triplets of submatrices, derived from two replicates and one non-replicate from a given locus (**Extended Data S2a**). Initially, following HiCRep [23], we smoothed all our input contact maps using a 2D mean filter with a smoothing factor *h* = 11 for Hi-C at 25 kb resolution. This smoothing filter replaces each entry in the contact map with the average counts of all contacts in its size-*h* neighborhood. We then identify, for a given biosample, one BR pair and a corresponding NR experiment. We randomly sample a 224 × 224 diagonal submatrix from the first biological replicate as our anchor input. The corresponding submatrix from the second biological replicate serves as the positive input, and the corresponding matrix from the non-replicate serves as the negative input.

For each example in the mini-batch, we begin by randomly selecting one BR Hi-C pair out of the 24 available human BR pairs. From this selected pair, we then randomly choose a pair of 224 × 224 diagonal submatrices from the same genomic region within the BR Hi-C pair. This process is repeated 256 times to from a set of 256 anchor-positive submatrix pairs. Next, for every BR submatrix pair in the set, we sample a submatrix from the same genomic region, but from the NR Hi-C experiments, as the negative submatrix to form a triplet for training. However, because there many NR pairs for each BR pair, it is impractical to sample all possible triplets. Instead, for every anchor-positive submatrix pair we sample 10 NR submatrices. This results in a total of 2,560 triplets within the mini-batch for training purposes. For one training epoch, we include 1000 iterations, with each iteration involving training on these 2,560 triplets.

The model employs a conjoined network architecture, wherein the three inputs are fed in parallel into the encoder, resulting in three embeddings: the anchor embedding *a*, the positive embedding *p*, and the negative embedding *n*. We aim for *a* and *p* to be as similar as possible, while *a* and *n* should be as dissimilar as possible. To achieve this, we use the triplet margin loss:

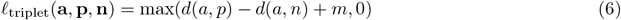

where *d* represents the cosine distance between two embeddings, *d*(*a, p*) = 1 − *cosine*(*a, p*), and *m* is a margin. We use *m* = 1 in our experiments.

Similar to the pre-training phase, the model is optimized using the AdamW optimizer with a cosine decay learning rate schedule. We use an initial learning rate of 10^−3^, weight decay of 0.05, a batch size of 256,and momentum parameters of *β*1 = 0.9 and *β*2 = 0.95. The fine-tuning consists of 50 epochs, with 5 epochs designated as warm-up epochs [85].

Following the fine-tuning process, we employ the trained model by scanning along the diagonal of a given Hi-C matrix with a submatrix size of 224 × 224 and a step size of 100. The overall reproducibility score between two Hi-C matrices is calculated by averaging the cosine similarities of the embeddings of this series of diagonal submatrices. Similar to HiCRep and GenomeDISCO, the average is calculated separately for each chromosome (excluding chrM and chrY) and then averaged across different chromosomes.

#### Other methods

HiCRep [23] assesses the reproducibility of Hi-C data using a stratum-adjusted correlation coe”cient. We used the Python implementation [87] available at https://github.com/dejunlin/hicrep. Genome-DISCO [27] uses random walks to smooth the contact map and then computes a concordance score to compare the contact maps (https://github.com/kundajelab/genomedisco). HiC-Spector [28] uses spectral decomposition to quantify the reproducibility of the Hi-C maps (https://github.com/gersteinlab/HiC-spector).

### Fine-tuning HiCFoundation for the loop detection task

To train HiCFoundation for loop detection, we use HiCCUPs [71] to identify loops that are observed in two BR Hi-C experiments. We begin by independently calling loops from two BR Hi-C matrices at 10 kb resolution, restricting to regions within 5 Mb of the diagonal. We use the recommended settings for HiCCUPs [71], including a peak width of 2, window size of 5, cluster radius of 20 kb, and false discovery rate (FDR) threshold of 0.1. We then identified consensus loop calls from two BRs (with centers ≤ 50 kb apart [21]) as our target for training. We define the neighboring 5 × 5 pixels (50 kb × 50 kb) of the loop calls as loop regions *L*. Next, we identify unlabeled regions by running HiCCUPs with a relaxed FDR threshold of 0.35. We gather all loop calls from any BR with this relaxed setting, excluding those already included in the consensus loops obtained earlier. The neighboring 5 × 5 pixels of these excluded loops are designated as unlabeled regions *U*. All remaining pixels are assigned as background regions *B*.

After assigning a label to each pixel of the Hi-C contact matrix, we randomly sample two 224 × 224 submatrices from the BR1 and BR2 Hi-C contact maps that include consensus loops from *L*. We then combine the two submatrices using a coe”cient *α* to create a mixed submatrix as the input for the model. Here, *α* is uniformly sampled from [0, 0.2, 0.4, 0.6, 0.8, 1.0]. We apply this data augmentation step to ensure that the consensus loop is detectable from any mixed submatrix. The corresponding pixel assignments from *L, U*, and *B* are sampled as the target *G*. Mini-batches for training the model are generated by randomly sampling such submatrices and corresponding target submatrix at different genomic regions from different Hi-C experiments.

The HiCFoundation model processes a submatrix as input and generates a corresponding pixel-level loop prediction matrix *P*. The entire framework is then optimized using the Dice loss [88] (**Extended Data S2b**),

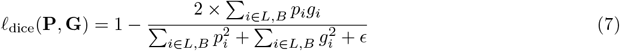

where 𝓁 _dice_ represents the Dice loss of a pixel-wise loop prediction matrix *P* and a corresponding ground truth loop detection *G, p*_*i*_ ∈ *P* is the predicted probability of the *i*th pixel in the prediction matrix, and *g*_*i*_ ∈ *G* is the ground truth loop assignment of the *i*th pixel. We only consider consensus loop regions and background regions (*i* ∈ *L, B*) during optimization, and any pixels belonging to the unlabeled regions *U* are not considered. The smoothing factor *ϵ* is set to 1 10^−6^. With the defined Dice loss to optimize the loop detection task, all other fine-tuning settings remain the same as those used for the reproducibility analysis. In particular, we freeze the pre-trained encoder and initialize the decoder weights using the the pre-trained decoder.

After fine-tuning, we used the fine-tuned model to detect loops by scanning the merged Hi-C contact map using a submatrix size of 224 × 224 and a step size of 100 at 10 kb resolution. The final prediction for a given pixel is the mean of the predictions from each overlapping submatrix. This process enabled us to obtain pixel-wise loop probability predictions for the input Hi-C contact map within the 5 Mb off-diagonal region.

To obtain the final loop calls from pixel-wise predictions, we use the mean-shift algorithm [39] to cluster pixels into loop calls. The mean-shift algorithm is a non-parametric clustering algorithm widely used for image processing and analysis. In our setting, the mean-shift algorithm takes a pixel-wise prediction matrix *P* as input, where each pixel value *P*_*i*_ (computed loop probability values, in our case) has corresponding 2D coordinates *x*_*i*_. The algorithm iteratively updates the matrix values and the corresponding coordinates. Here we only consider points with confident predictions (*P*_*i*_ ≃ 0.9) from HiCFoundation. The coordinates of pixel *i* are iteratively updated following 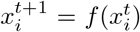 until convergence when 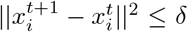, with *δ* set to 0.001. Here the update function is

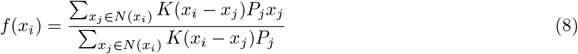

where *N* (*x*_*i*_) is the set of neighboring grid points of *x*_*i*_, with locations satisfying ||*x*_*j*_ −*x*_*i*_ ||^2^ ≤2*σ*, and *K*(*p*) is a Gaussian kernel function with bandwidth *σ* = 2:

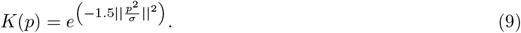

At the same time, the value of pixel *i* is also updated by

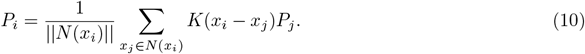

After applying the mean-shift algorithm, all points tend to move toward nearby locations with the highest density. Subsequently, any shifted points that are closer than a threshold distance of 2 are clustered together, and the grid point with the maximum density or probability is selected as the final loop detection point.

We also fine-tune HiCFoundation for loop calling using LC Hi-C data. To simulate LC Hi-C, we down-sample the Hi-C contact map to 1/16 of the original read count. Specifically, for a Hi-C contact map with a total read count *K*, the probability of sampling a read count *k* is 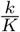. This process is repeated 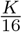 times without replacement. Keeping all other settings fixed, we fine-tune HiCFoundation with the downsampled Hi-C data as input and the labels derived from the original data. This process results in a new model tailored specifically for LC Hi-C data.

#### Performance evaluation

To evaluate loop detection performance, we use the F1 score, computed by comparing the loop predictions against the consensus ground truth loops. The loop F1 is defined as

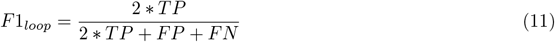

where TP represents true positive loop calls, which are predicted loop calls within a 5-pixel range of a ground truth loop; FP stands for false positive loop calls that only appear in the predicted loops; and FN indicates false negative loops that are only called in the ground truth but are not detected by the predictions with the specified range tolerance.

#### Parameter settings of comparison methods

For HiCCUPs, we use juicer tools.jar (v1.22.01) downloaded from https://github.com/aidenlab/juicer/wiki/Download. The command for general loop calling is `java -Xmx64g -jar juicer tools.jar hiccups -r 10000 -f 0.1 -p 2 -w 5 -d 20000 -k KR {input.HiC} {output loop.bedpe}`. For relaxed loop calls, the FDR threshold (-f) is changed from 0.1 to 0.35. For Chromosight, we downloaded and installed the code from https://github.com/koszullab/chromosight. The command for the loop calling is `chromosight detect --thread 8 --max-dist 5000000 --pattern=loops small {input.cool} {output prefix}`. The loop calls are saved in output prefix.tsv. For Mustache, we downloaded and installed the code from https://github.com/ay-lab/mustache. The command for loop calling is `mustache -f input.hic -r 10000 -o output.loop -p 16 -d 5000000`, where “-p” specifies the number of parallel processes to run.

### Fine-tuning HiCFoundation for the resolution enhancement task

For the resolution enhancement task, we adopt the same network architecture as in previous tasks. During the fine-tuning stage, we transfer the initial weights from the final pre-trained model, freeze the encoder weights, and only fine-tune the decoder weights.

To train the HiCFoundation model for resolution enhancement, we prepare the LC Hi-C submatrices as input and the HC Hi-C submatrices as output. As in previous work [33, 41, 74, 21], we downsampled the Hi-C contact map to 1/16 of the original read count to obtain a LC matrix for each HC Hi-C matrix This downsampling is done via iterative sampling. For a Hi-C contact map with a total read count *K*, the probability of sampling a read at a locus pair with *k* reads is 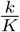. This sampling operation is repeated 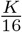 times without replacement to obtain a LC Hi-C matrix with 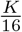 total reads. Furthermore, similar to most previous resolution enhancement methods, we carry out the analysis using 10 kb resolution, and we restrict the model to consider only intra-chromosomal contacts that span up to 2 Mb. Note that, given HiCFoundation’s input submatrix size of 224 ×224, our model covers a slightly larger region of 2.24 Mb. We randomly sample 224 ×224 diagonal submatrices from the down-sampled LC Hi-C *X* as our input and the corresponding diagonal submatrices from HC Hi-C as our target *Y*. Considering the limited biological significance of diagonal counts, we set the diagonal reads in each submatrix to 0. We further normalize the input *X* and target *Y* for better optimization. Specifically, the HC target *Y* is clamped to [0, 1000] and then normalized to [0, 1] using min-max normalization. Similarly, the LC input *X* is clamped to [0, 100] and normalized to [0, 1] using min-max normalization.

Taking the LC submatrix *X* as input, the HiCFoundation model generates a corresponding enhanced submatrix *P*. With the HC submatrix *Y* as the target, the entire framework is optimized via pixel-wise MSE:

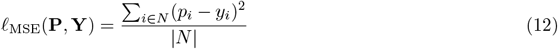

where *N* is the set of all non-diagonal pixels in the submatrices, *p*_*i*_ is the *i*th pixel’s prediction by HiCFoundation, and *y*_*i*_ is the *i*th pixel’s target from *Y*. All other fine-tuning settings remain the same as those used for the reproducibility analysis and loop detection tasks.

After fine-tuning, we use the fine-tuned model to enhance the LC Hi-C data. This enhancement is achieved by scanning the LC Hi-C contact map using a submatrix size of 224 ×224 and a small step size of 20 along the diagonal at a resolution of 10 kb to ensure that all off-diagonal 2 Mb regions are enhanced. The final enhancement output for a given pixel is determined by averaging the predictions from each overlapping submatrix. This process allows us to obtain pixel-wise enhancement results for the input LC Hi-C contact map within the 2 Mb off-diagonal region.

#### Performance evaluation

To evaluate resolution enhancement performance, we followed the comprehensive evaluation pipeline in [42]. Specifically, the evaluation includes six metrics:

1. **Pearson’s correlation coeffcient (PCC)**: Considering the genomic distance effect in Hi-C matrices, the correlation measurement is not performed across the Hi-C matrix. Instead, the correlation is assessed by comparing corresponding diagonals of the two input matrices. Specifically, the correlation score is defined as

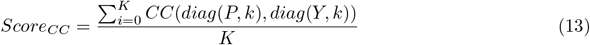

where *Score*_*CC*_ is the correlation coe”cient score; *CC* is the Pearson correlation coe”cient function; *K* is the number of diagonals considered, set to 100; *diag*(*P, k*) refers to the *k*-th diagonal above the main diagonal from the enhanced Hi-C matrix *P* ; and *diag*(*Y, k*) refers to the *k*-th diagonal above the main diagonal from the HC Hi-C matrix *Y*.
2. **Spearman’s correlation coeffcient (SCC)**: Same as PCC, but using the Spearman correlation coe”cient for *Score*_*CC*_.
3. **Structural similarity index measure(SSIM)**: Defined in Eq.4, calculated with respect to the enhanced and HC Hi-C matrices.
4. **Peak signal-to-noise ratio (PSNR)**: PSNR is a ratio between the maximum possible value (power) of a signal and the power of distorting noise that affects the quality of its representation:

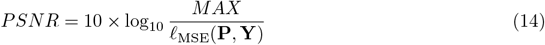

where *MAX* is the maximum signal value, which is 1 because of our normalization; 𝓁_MSE_(**P, Y**) measures the MSE (defined in Eq. 12) between the enhanced matrix *P* and HC matrix *Y*.
5. **Loop F1 score (F1**_*loop*_): For loop detection, we employed the fine-tuned loop HiCFoundation model as our loop caller. The loop F1 score (Eq. 11) is evaluated by comparing the loop calling results from the enhanced Hi-C data with the loop calls derived from the HC Hi-C data.
6. **TAD F1 score (F1**_*T AD*_): Similarly, the TAD F1 score is computed using the Insulation Score [89] for TAD detection, as implemented in [21], applied separately to the enhanced and HC Hi-C data.

The calculation of these six metrics is done at the chromosome level instead of the submatrix level. The final scores are averaged across chromosomes. We used the mean rank as our final score, which is defined as the mean of the ranks from the different methods on the different metrics:

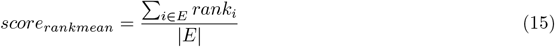

where *E* is the set of evaluation metrics, and *rank*_*i*_ is the rank of *i*th metric from the three evaluation categories. Thus, a smaller mean rank is better.

#### Comparisons with other methods

To compare against other methods, we follow the training code and default training settings for each method, using our training data and benchmarked on our test set. The HiCSR, HiCNN, and HiCARN code are from, respectively, https://github.com/PSI-Lab/HiCSR, http://dna.cs.miami.edu/HiCNN2, and https://github.com/OluwadareLab/HiCARN.

### Benchmarking on the multi-species dataset

To benchmark HiCFoundation on multi-species dataset, we collected 337 Hi-C experiments from the DNA Zoo database (https://www.dnazoo.org/).

For the HiCFoundation-Reso benchmark, we downsample the Hi-C contact maps to 1/16 of their original read count to generate LC Hi-C inputs for HiCFoundation-Reso. Performance is then assessed by comparing the enhanced Hi-C data against the raw Hi-C data using six previously defined evaluation metrics, along with the overall summary mean rank.

To investigate the relationship between performance and species similarity relative to humans, we use TimeTree [90] to measure species divergence time from humans. Only human data are used to train different HiCFoundation downstream models.

### Fine-tuning HiCFoundation for the epigenomic signal prediction task

For the epigenomic signal prediction task, we adopt the same network architecture as in the other HiCFoundation tasks, with model weights initialized using the final pre-trained model. The HiCFoundation-epi model takes a Hi-C submatrix as input and outputs six different epigenomic signals. Accordingly, to map the output embedding of HiCFoundation-epi to various track signals, six distinct, fully connected layers are used, each processing the embedding independently. During the fine-tuning stage, the encoder is frozen, and the decoder and fully connected layers are updated using objective defined below.

To train the HiCFoundation-epi model, we first process the Hi-C and other epigenomic signal data to a resolution of 1 kb. For the input Hi-C data, we use the total interaction counts binned at 1 kb resolution as input. For the output epigenomic signal, we use the normalized read-depth signal, as provided in the files from ENCODE and 4DN, and then average these at the same 1 kb resolution as the target. We further normalize each epigenomic signal track for better optimization by first clamping to the 98th percentile (non-zero) value and then normalizing to [0, 1] using min-max normalization. We generate training examples by scanning the Hi-C matrix along the diagonal with 128 ×4000 submatrix, with a stride of 64. Each such submatrix thus represents the entire 2 Mb diagonal region associated with a specific bin. We randomly sample batches of these submatrices at every training epoch. The position embedding in Equation 2 is applied to each submatrix, after adjusting for the size of the input. The corresponding epigenomic signals at each locus comprise the target output *Y*, with dimension 6 ×128. Taking the Hi-C submatrix *X* as input, the HiCFoundation model generates a corresponding predicted epigenomic signal matrix *P*.

With the experimental epigenomic track *Y* as the target, the entire framework is optimized via a loss function consisting of two terms. The first term is the mean-squared error loss between the predicted signal and the target experimental signal:

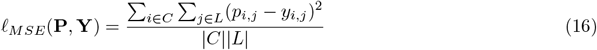

where *p*_*i,j*_ and *y*_*i,j*_ are, respectively, the predicted and target signals of the *i*th epigenomic track at the *j*th position in the ouput window *L*. The second term is the cosine loss between the predicted signal and the target experimental signal,

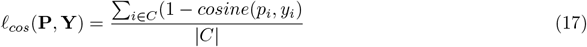

where *p*_*i*_ ∈ℝ^*L*^ and *y*_*i*_ ∈ℝ^*L*^ are, respectively, the predicted and target signals of the *i*th epigenomic track in a window of size *L*; *cosine*(*p*_*i*_, *y*_*i*_) measures the similarity between the predicted signal *p*_*i*_ and target signal *y*_*i*_. The combined loss function is

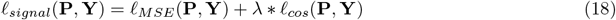

where *P* ∈ℝ^*C,L*^ and *Y* ∈ℝ^*C,L*^ are the predicted signal and target signals, respectively. Here, *C* = 6 is the number of epigenomic signal tracks, and *L* = 128 is the length of the signal window. The coe”cient *lambda*, which balances the two loss terms, is set to 1. During the optimization process, 𝓁_*MSE*_(**P, Y**) encourages locally accurate signal prediction, and 𝓁_*cos*_(**P, Y**) enforces the model to capture the global profiles of the target signal [91].

After the fine-tuning process, the model infers epigenomic data from Hi-C input by scanning the raw Hi-C contact map using a submatrix size of 128 ×2000 and a step size of 64 along the diagonal at a resolution of 1 kb. The overlapping predictions are averaged to obtain the final predictions. After this process, the predicted epigenomic tracks at 1 kb resolution are available for downstream analysis.

To evaluate the performance of HiCFoundation-epi, we compute the Pearson’s correlation between the predicted and target signal.

We include two baseline methods for comparison with HiCFoundation-epi. The first baseline is principal component analysis (PCA) of Hi-C data [2], implemented in Homer [92]. PCA is a widely used method for dimensionality reduction, which redefines a given coordinate system so that the data can be described with as few dimensions as possible. The axes of this new coordinate system are called “principal components.” The first principal component is identified to account for the maximum variance in the data, while the second principal component captures as much of the remaining variance as possible, and so on. PCA is often applied to full chromosome matrices to identify important features, with the principal component serving as a 1D view of the Hi-C matrix that can be compared with different tracks. The second baseline is a network with the same architecture as HiCFoundation-epi, but without the pre-training stage. We adopted the same input, output, and optimization settings as HiCFoundation, but instead of transferring weights from pre-training and using a frozen encoder, the encoder and decoder are randomly initialized and optimized by the same loss objectives used in HiCFoundation. Because the encoder is optimized during training, this approach requires extra backpropagation through the encoder, resulting in a training time approximately five times longer than the fine-tuning of HiCFoundation.

### Fine-tuning HiCFoundation for single-cell resolution enhancement

For single-cell Hi-C analysis, we adopt the same fine-tuning setup as we do for bulk Hi-C analysis. We focus on the single-cell Hi-C resolution enhancement task, where the model takes low-coverage scHi-C as input and outputs corresponding high-coverage scHi-C at the same resolution.

We prepared the training examples in three steps. First, each raw scHi-C matrix is binned at a resolution of 1MB. Second, for each chromosome, the raw matrix is either padded or randomly cropped to a fixed size of 224 ×224 to ensure compatibility with the input dimensions of HiCFoundation. Third, a downsampled matrix is generated by randomly reducing the number of contacts by a specified proportion (1/4). The downsampling procedure is the same as in the resolution enhancement task for bulk Hi-C.

For training the model, the downsampled scHi-C matrix is used as the input, while the raw scHi-C matrix serves as the target for fine-tuning HiCFoundation. We use a weighted MSE loss function, in which the weighting mechanism aims to reduces the sensitivity of the loss gradients to large values, defined as

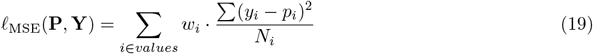

where *values* is the collection of distinct true values at each batch training, *i* is any value in *values, w*_*i*_ is the weight for the MSE loss on value *i, y*_*i*_ represents all the true values of *i, p*_*i*_ are the predicted values of *i*, and *N*_*i*_ is the number of true value *i*. The weights range from 0 to 1, increasing linearly with the rank of the true value:

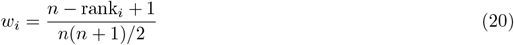

where *n* is the total number of distinct values, and *rank*_*i*_ is the rank position of *i* in ascending order.

The model, HiCFoundation-sc, is fine-tuned for 50 epochs, updating only the weights in the decoder. We use the AdamW optimizer with parameters of *β*_1_ = 0.9 and *β*_2_ = 0.95. A warm-up learning rate adjustment strategy is applied, where the learning rate begins at 0 and gradually increases. After reaching a fixed learning rate of 0.003, it then decays linearly to zero at the last epoch.

For evaluation, we adopt five performance measures. This includes four of the measures used in the bulk Hi-C resolution enhancement task, as well as HiCRep [23], which was used in previous work [62]. Additionally, following Higashi [62], we applied quantile normalization to both the ground-truth and enhanced Hi-C matrices prior to calculating Pearson, Spearman and HiCRep.

#### Comparisons with other methods

To compare against other methods, we follow the training code and default training settings for each method, using our data and benchmarked on our test set. The scHiCluster, Higashi, and scVI-3D code are from, respectively, https://github.com/zhoujt1994/scHiCluster, https://github.com/ma-compbio/Higashi, and https://github.com/yezhengSTAT/scVI-3D.

**Extended Data S1:**
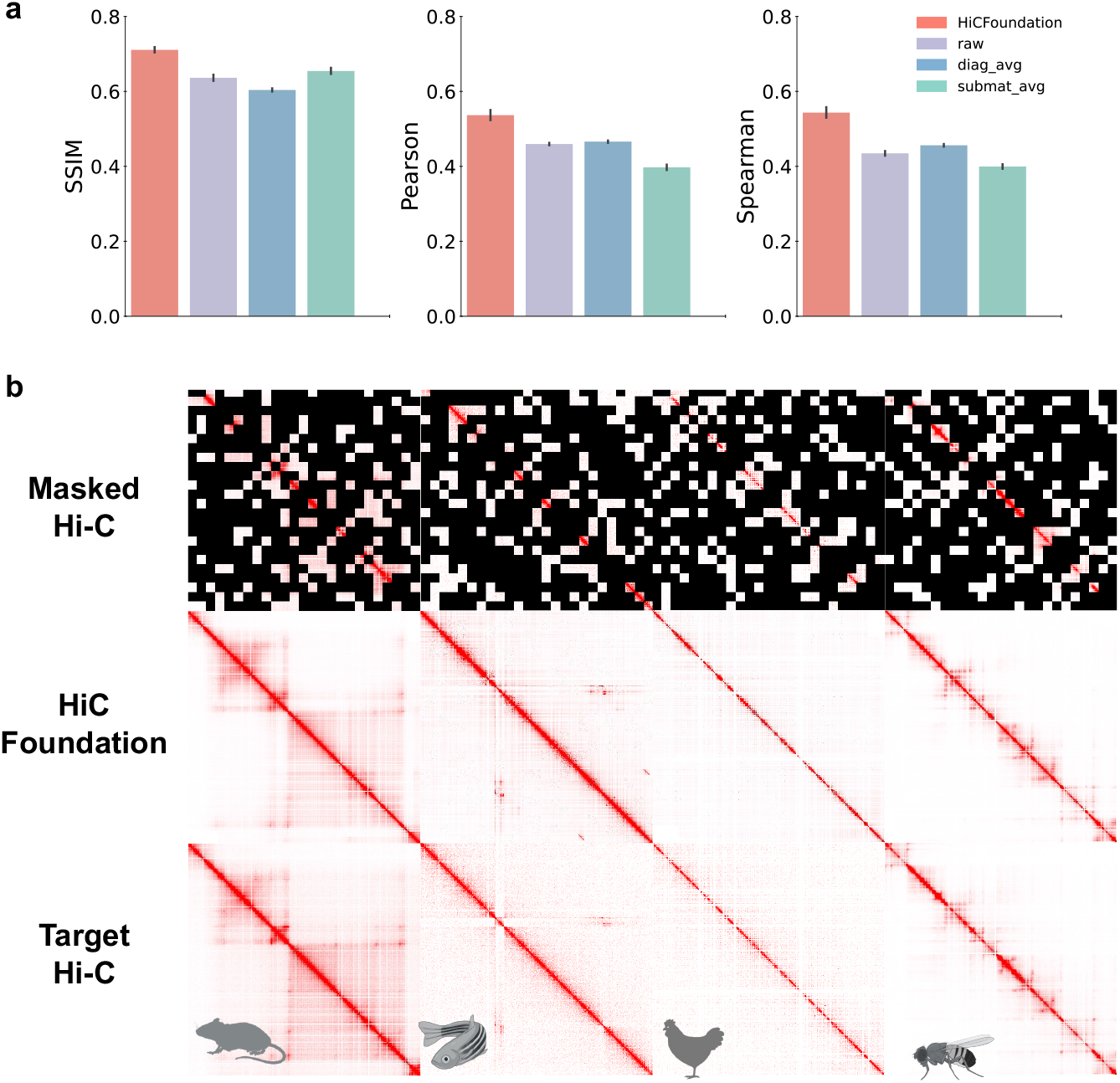
Benchmark of pre-training performance. **a**. Performance comparison of different methods on reconstruction. **b**. Visual example of HiCFoundation on testing sets across different species.

**Extended Data S2:**
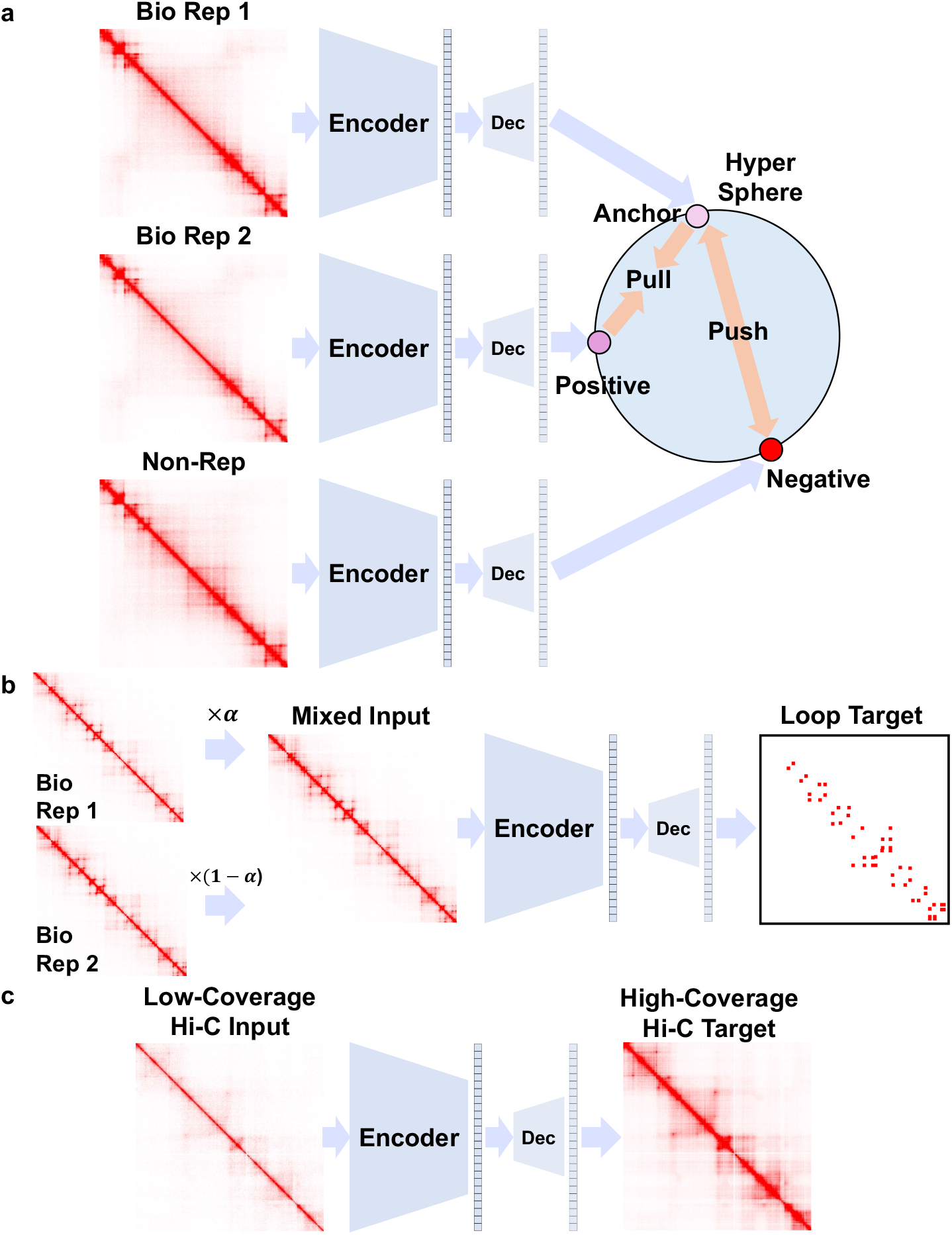
Fine-tune protocol of HiCFoundation for different Hi-C analysis tasks. During fine-tuning, the encoder remains frozen and the decoder is updated according to task-specific signals. **a**. Fine-tuning protocol for reproducibility analysis. The model takes as input submatrices from biological replicate 1, biological replicate 2, and a randomly sampled non-replicate, generating corresponding embeddings. The framework is then optimized using a triplet loss function, which minimizes the distance between embeddings of the biological replicates while maximizing the distance between the embeddings of the biological replicate and the non-replicate. **b**. Fine-tuning protocol for the chromatin loop detection task. During fine-tuning, corresponding submatrices from two BR experiments serve as the input, and consensus loops from HiCCUPs are used as the target for optimizing the decoder. **c**. Fine-tune protocol for resolution enhancement. The decoder is optimized to generate high-coverage Hi-C by taking the low-coverage Hi-C as input.

**Extended Data S3:**
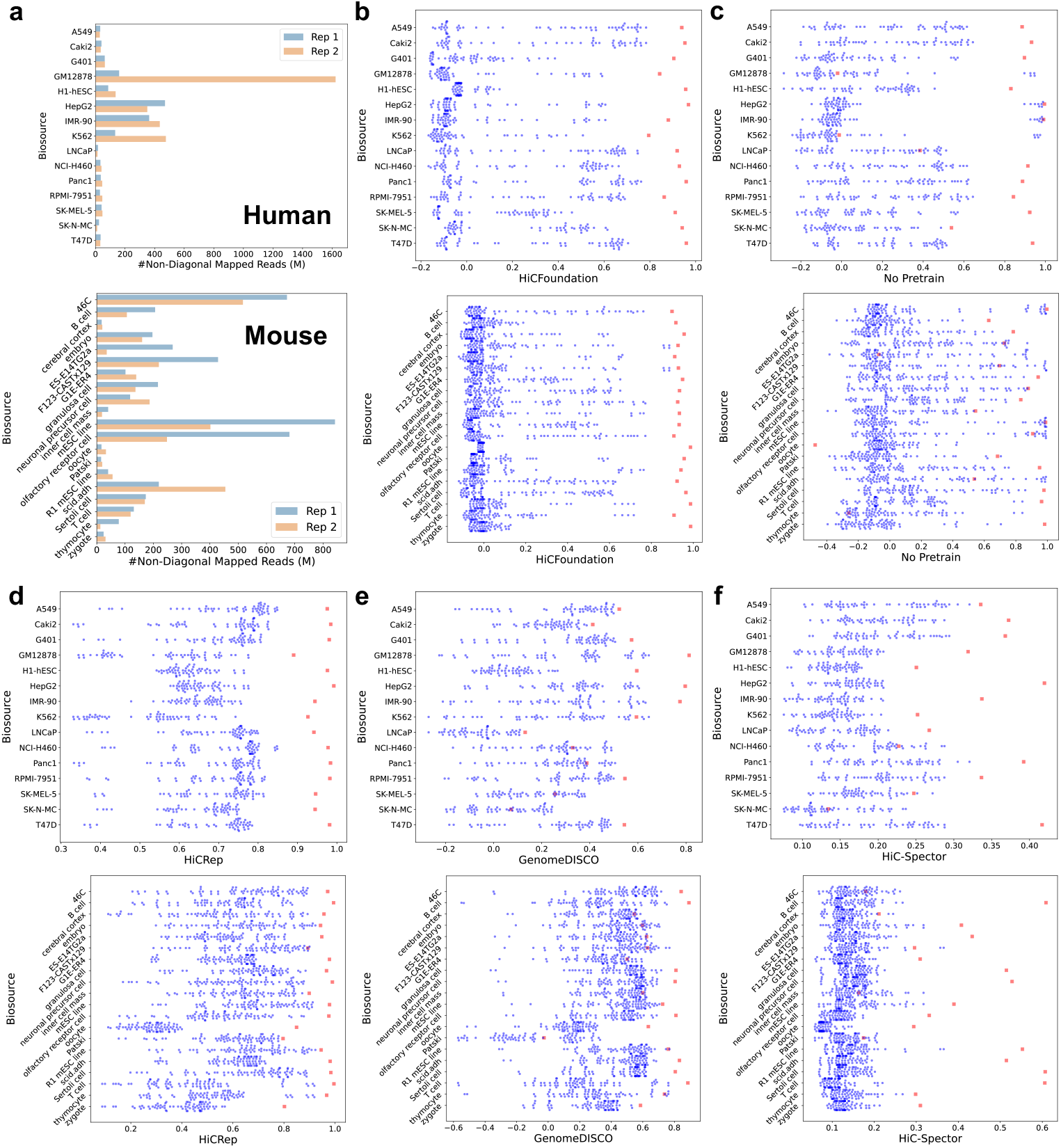
Reproducibility Analysis of other methods. **a**. The total number of non-diagonal mapped reads of biological replicates in the reproducibility testing dataset, including human and mouse data. **b**. Reproducibility scores of HiCFoundation on different cell types from the human and mouse datasets, respectively. Here a red square indicates the score between biological replicates and the blue dots refers to the non-replicates. **c**. Reproducibility scores of the No-pretrain model on different cell types from human and mouse datasets. **d**. Reproducibility scores of HiCRep on different cell types from human and mouse datasets. **e**. Reproducibility scores of Genome-DISCO on different cell types from human and mouse datasets. **f**. Reproducibility scores of HiC-Spector on different cell types from human and mouse datasets.

**Extended Data S4:**
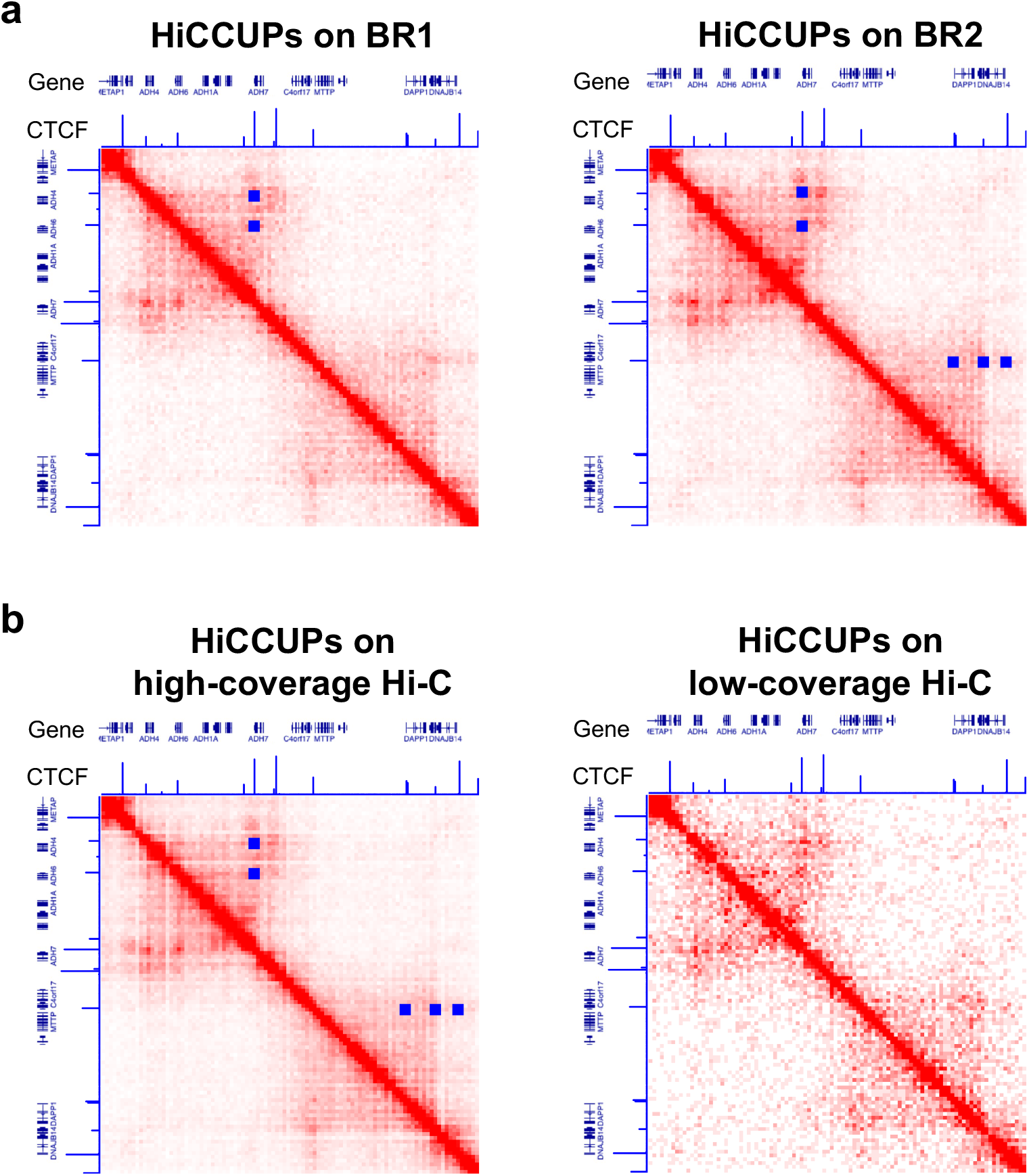
Loop detection example by HiCCUPs. The panels include Hi-C data, corresponding CTCF ChIP-seq signals (ENCSR000EFI), reference genes, and loop detections (blue squares). **a**. HiCCUPs loop call on two biological replicates. The consensus one are treated as our ground truth loop. **b**. HiCCUPs loop call on high-coverage Hi-C and the low-coverage Hi-C.

**Extended Data S5:**
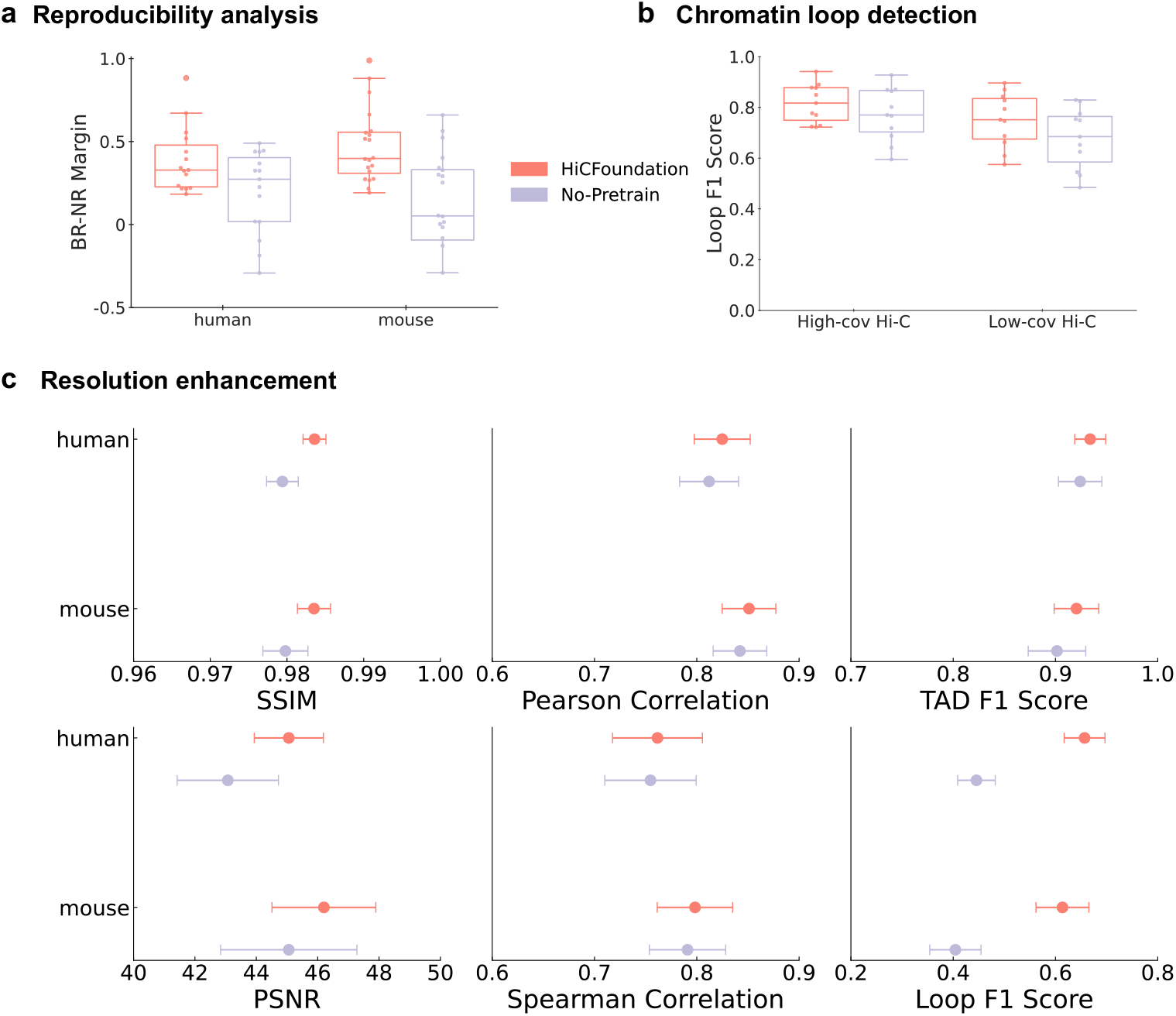
Benchmark of HiCFoundation and No-pretrain on different chromatin architecture analysis tasks. **a**. Comparisons on reproducibility analysis. **b**. Comparisons on chromatin loop detection. **c**. Comparisons on resolution enhancement.

**Extended Data S6:**
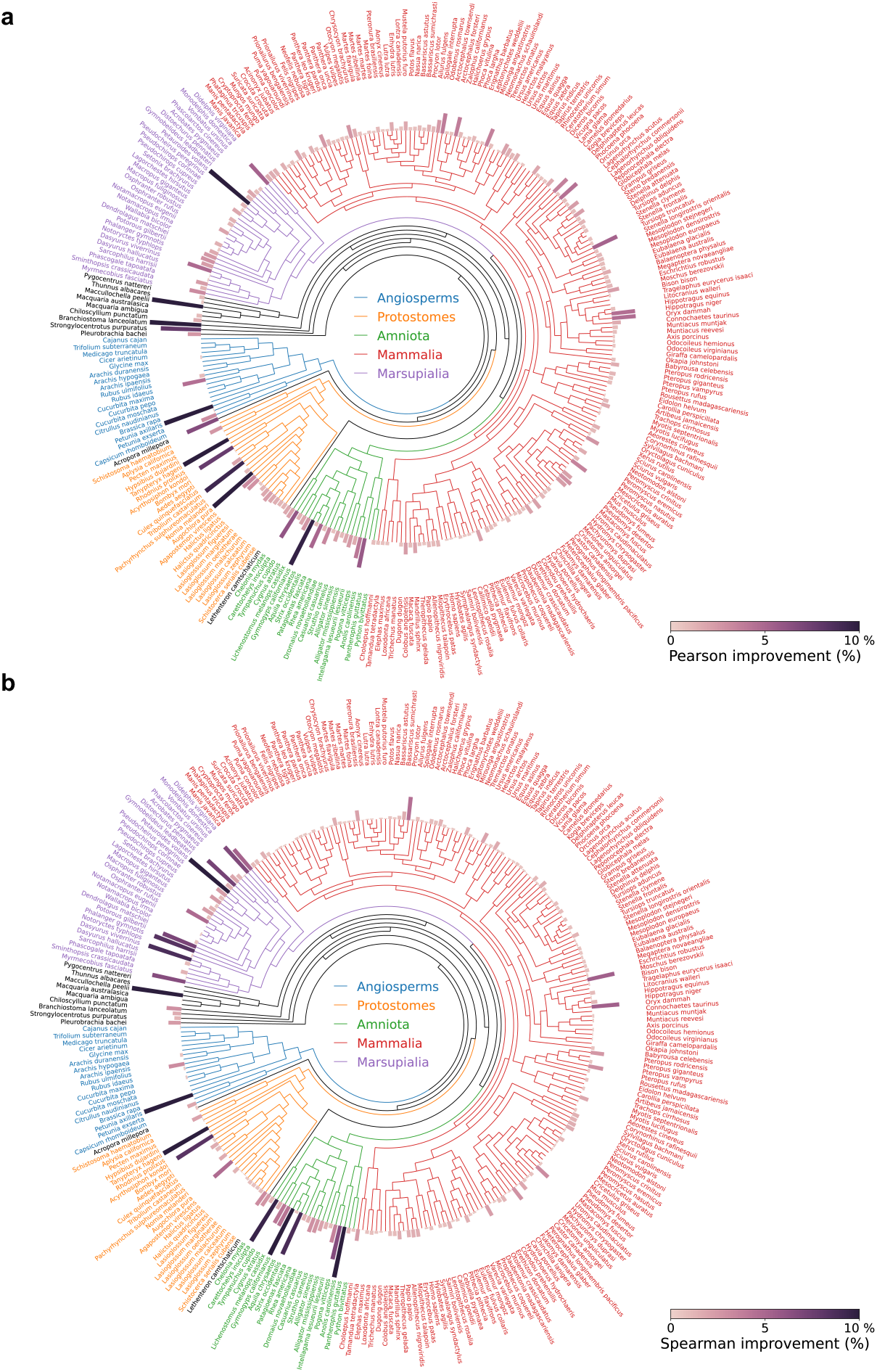
HiCFoundation improvement over No-pretrain on multi-species DNA Zoo dataset. **a**. Pearson correlation improvment of HiCFoundation relative to No-pretrain. **b**. Spearman correlation improvment of HiCFoundation relative to No-pretrain.

**Extended Data S7:**
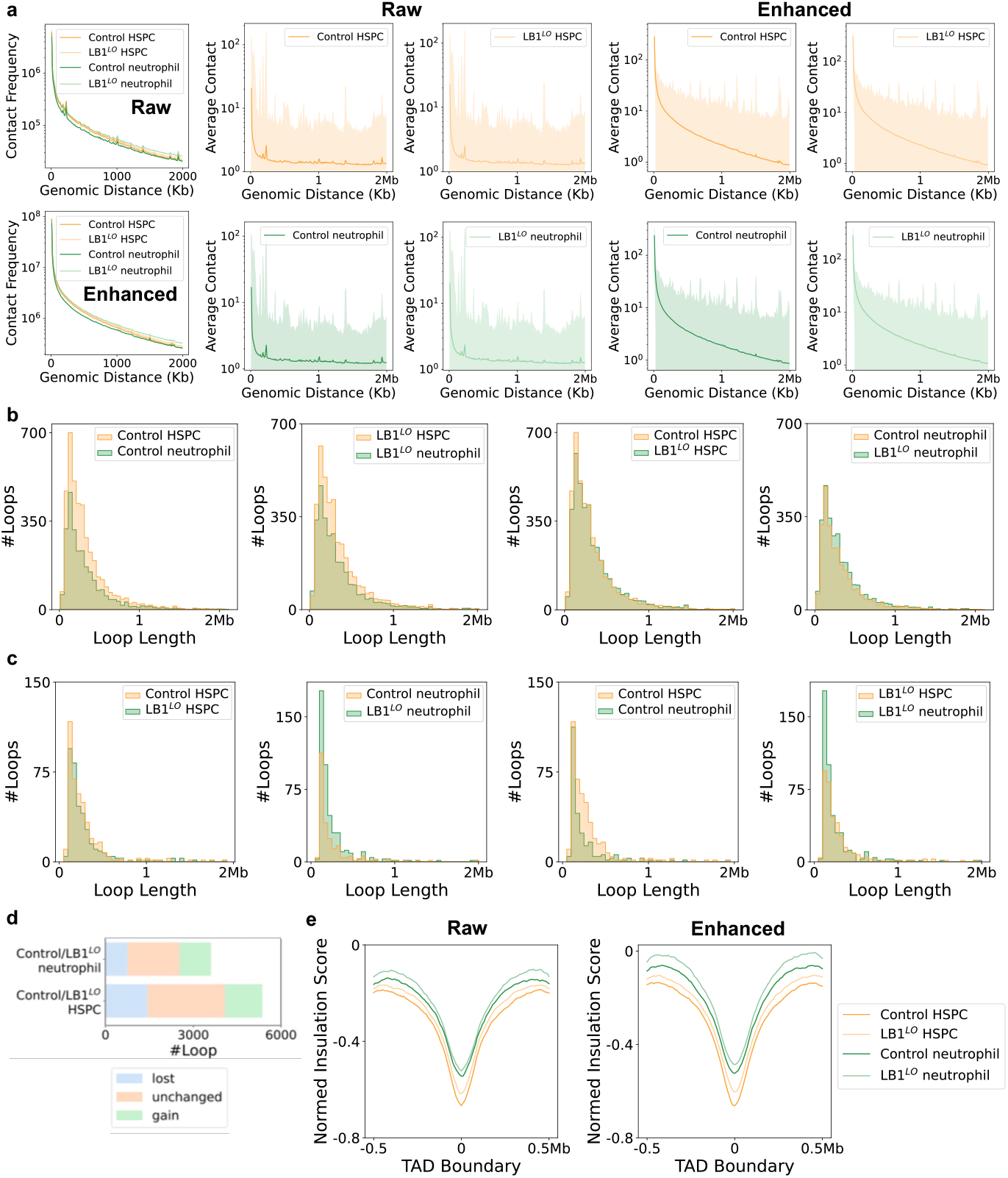
Chromatin architecture change analysis in HSPCs and neutrophils. The comparisons are made from four different samples: control HSPC, LB1^*LO*^ HSPC, control neutrophil, and LB1^*LO*^ neutrophil. Here LB1^*LO*^ represents the loss of lamin B1, **a**. Contact frequency as a function of genomic distance for different samples. The comparisons between and enhanced data are presented. The right panel provides a detailed view of the mean whole-genome Hi-C contact frequencies (normalized for depth) over genomic distance for different samples. **b**. Loop length comparison of different samples with enhanced Hi-C. **c**. Loop length comparison of different samples with raw Hi-C. **d**. Loop change comparison under two settings: control neutrophil vs. LB1^*LO*^ neutrophil, and ontrol neutrophil vs. LB1^*LO*^ HSPC. **e**. Average insulation score around TAD boundaries (±0.5 Mb). The two panel represents the raw and enhanced results, respectively.

**Extended Data S8:**
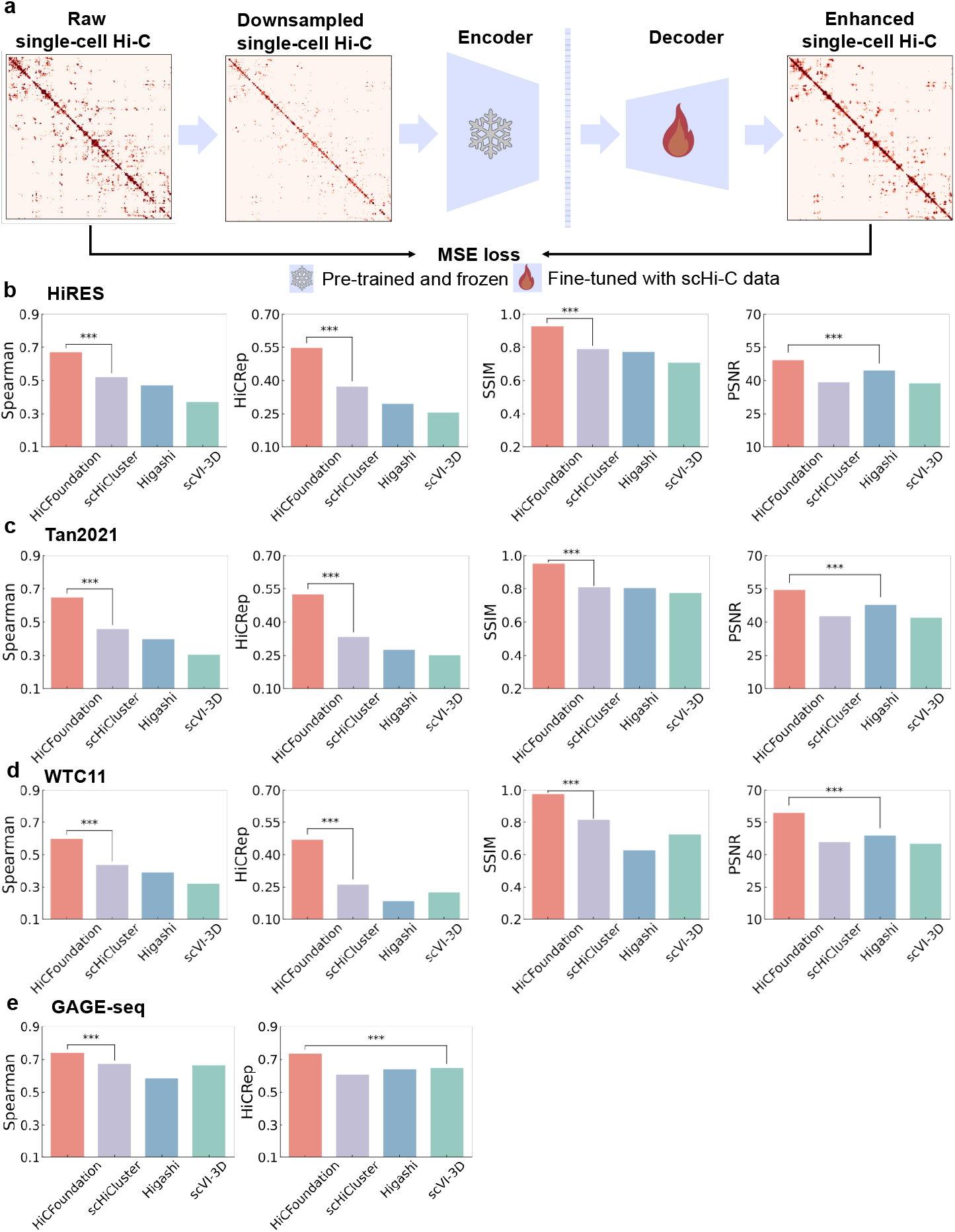
HiCFoundation for single-cell Hi-C analysis. **a**. The fine-tuning pipeline of HiCFoundation for single-cell Hi-C resolution enhancement. HiCFoundation-sc takes a downsampled scHi-C matrix as input and is fine-tuned to generate the original scHi-C matrix. **b**. Benchmark of HiCFoundation-sc on HiRES dataset, consisting of single cells from developing mouse embryos. **c**. Benchmark of HiCFoundation-sc on Tan2021 dataset, consisting of single cells from mouse forebrain cortex. **d**. Benchmark of HiCFoundation-sc on human WTC11 dataset. **e**. Benchmark of HiCFoundation-sc on GAGE-seq datasets from the K562 human cell line. Panels **b-e** include other evaluation metrics that are not listed in **Fig. 6b**.

**Extended Data S9:**
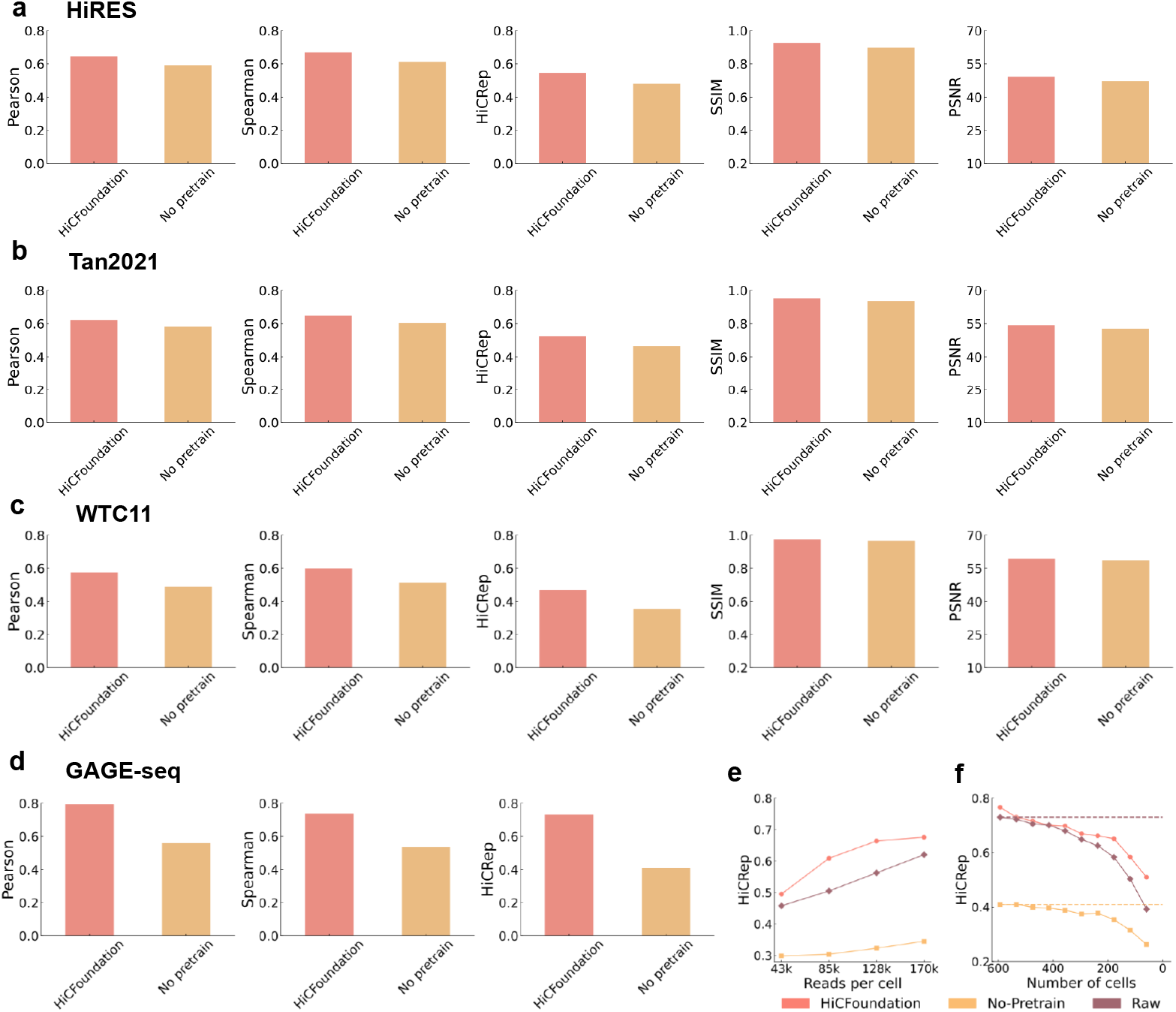
Comparison of HiCFoundation and No-pretrain for single-cell analysis. **a**. Benchmark of HiCFoundation-sc on HiRES dataset, consisting of single cells from developing mouse embryos. **b**. Benchmark of HiCFoundation-sc on Tan2021 dataset, consisting of single cells from mouse forebrain cortex. **c**. Benchmark of HiCFoundation-sc on human WTC11 dataset. **d**. Benchmark of HiCFoundation-sc on GAGE-seq datasets from K562 human cell line. **e**. Benchmark of HiCFoundation-sc under different reads per cell from the GAGE-seq dataset. The panel includes evaluation metrics that are not included in **Fig. 6e. f**. Benchmark of HiCFoundation-sc under different number of cells from the GAGE-seq dataset. The panel includes evaluation metrics that are not included in **Fig. 6f**.

## Notes

### Competing Interest Statement

The authors have declared no competing interest.

