## Supplementary_Note for "A generalizable Hi-C foundation model for chromatin architecture, single-cell and multi-omics analysis across species"

### Supplementary information

#### Supplementary Note 1: Inconsistency of Hi-C normalization during training and inference

In this work, we directly benchmarked resolution enhancement methods on raw Hi-C data, whereas previous works trained and tested their models on normalized Hi-C data [33, 41, 74, 21]. We work with raw rather than normalized data because the resulting model is more applicable to the realistic setting in which LC data needs to be enhanced.

To understand the problem with trying to enhance normalized data, consider the steps involved in training and then applying such a model (**Fig. S1a**). Initially, raw Hi-C data is normalized, typically using Knight-Ruiz normalization or matrix balancing, to produce a normalized Hi-C matrix, which serves as the training target. The normalized Hi-C is then downsampled to generate the normalized LC Hi-C, which serves as the training input. The model is optimized using these pairs of training inputs and training targets. After optimization, the trained model is deployed to take the normalized LC Hi-C as input and output the enhanced normalized Hi-C. However, the input used here does not correspond to the normalized LC input encountered in real testing scenarios, because that data was normalized using HC Hi-C data.

To illustrate the problem, we compared the performance of various trained models on test data normalized using the two different procedures. For this experiment, we use HiCARN1, HiCARN2, HiCNN and HiCSR models trained in the standard "early normalization" fashion, using training data that was first normalized and then downsampled. We then compare the performance of each of these models on two different tests sets: one that was normalized first and then downsampled, and a second that was downsampled and then normalized.

---

Supplementary Figure S1 *(following page)*: **Illustration of the normed resolution enhancement task during training and inference.**. **a.** The inconsistent resolution enhancement setting is widely adopted by previous approaches. Initially, raw Hi-C data is normalized to produce a normalized Hi-C matrix, which serves as the training target. The normalized Hi-C data is then downsampled to generate the training input. Similar pairs are used for testing benchmarks. However, in real testing scenarios, KR normalization must be applied to LC Hi-C data to construct the inference input. The trained model then takes the normalized LC Hi-C as input and outputs the enhanced HC Hi-C. **b.** Consistent resolution enhancement setting. To ensure consistency of inputs during training and evaluation, we implemented a revised processing pipeline. During training, raw Hi-C data is first downsampled to generate raw LC Hi-C. Both raw Hi-C and raw LC Hi-C are then KR normalized separately, resulting in normalized Hi-C and normalized LC Hi-C, which serve as the training target and testing targets, respectively.

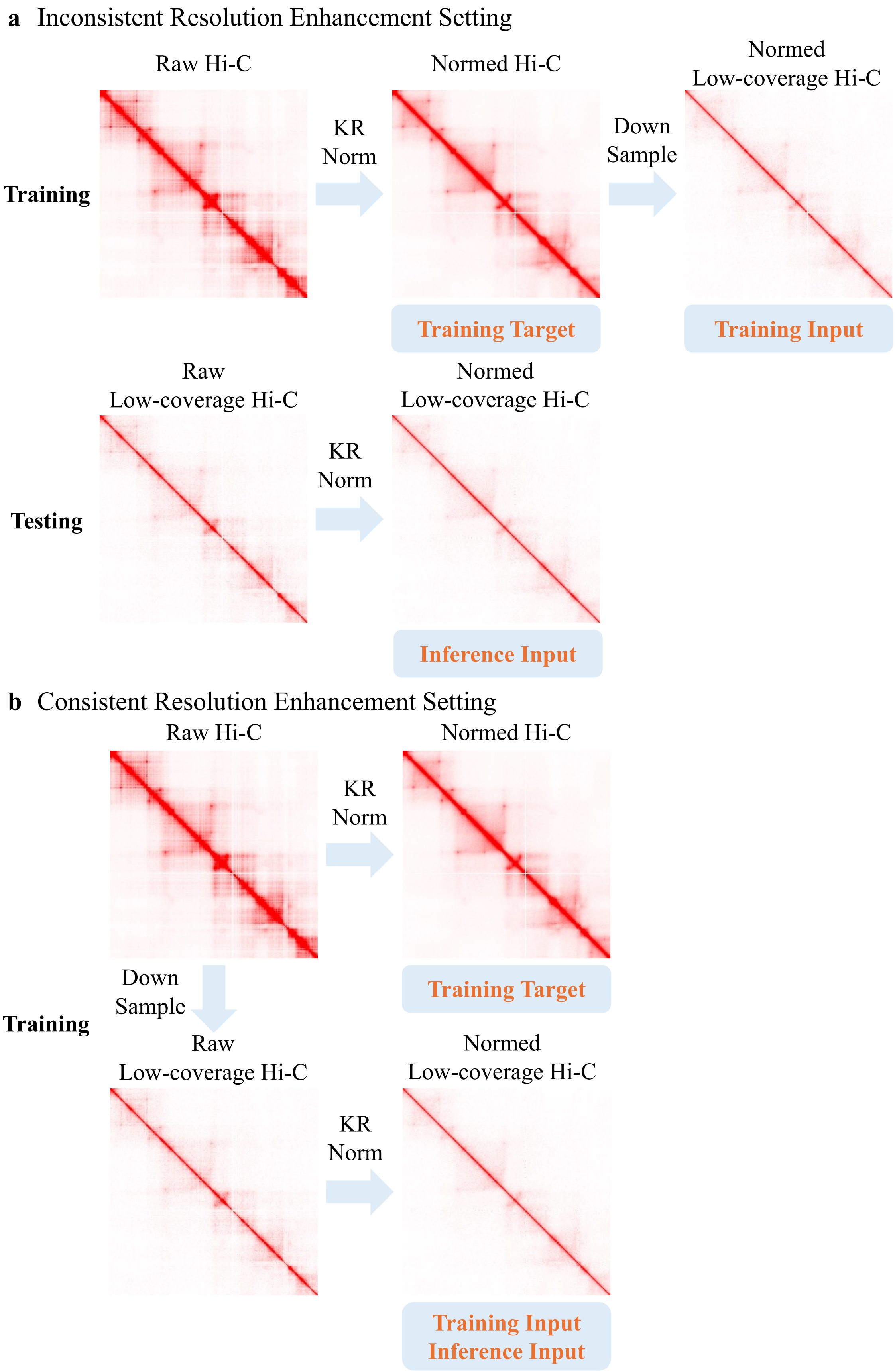
a Inconsistent Resolution Enhancement Setting
Raw Hi-C
Normed Hi-C
Normed
Low-coverage Hi-C
Training
KR
Norm
Down
Sample
Training Target
Training Input
Raw
Low-coverage Hi-C
Normed
Low-coverage Hi-C
Testing
KR
Norm
Inference Input
b Consistent Resolution Enhancement Setting
Raw Hi-C
Normed Hi-C
KR
Norm
Training Target
Down
Sample
Training
Raw
Low-coverage Hi-C
Normed
Low-coverage Hi-C
KR
Norm
Training Input
Inference Input

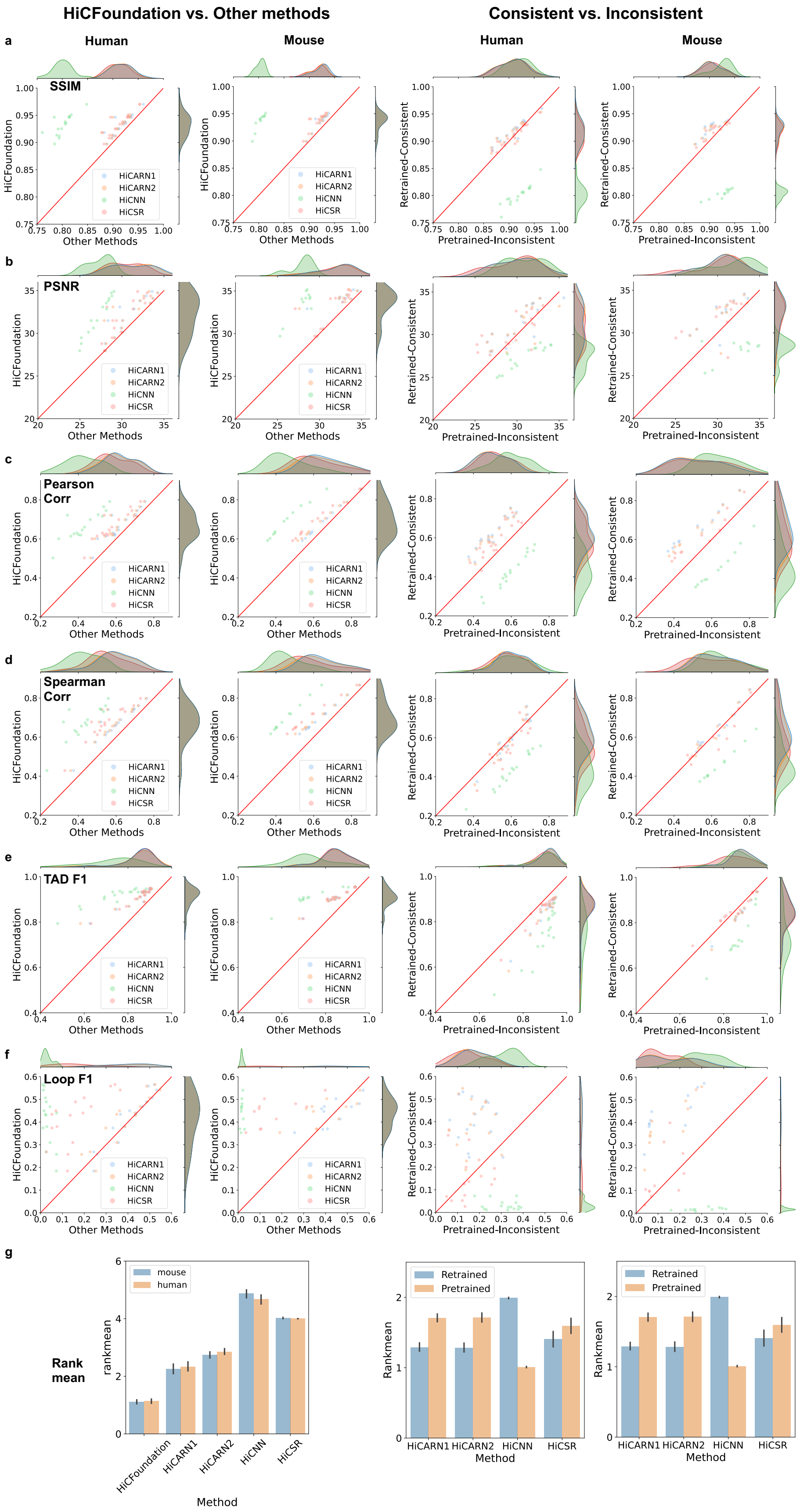
HiCFoundation vs. Other methods
Consistent vs. Inconsistent
a
Human
Mouse
Human
Mouse
SSIM
HiCFoundation
Other Methods
Retrained-Consistent
Pretrained-Inconsistent
HiCARN1
HiCARN2
HiCNN
HiCSR
b
PSNR
c
Pearson
Corr
d
Spearman
Corr
e
TAD F1
f
Loop F1
g
Rank
mean
rankmean
mouse
human
HiCFoundation
HiCARN1
HiCARN2
HiCNN
HiCSR
Method
Rankmean
Retrained
Pretrained

Supplementary Figure S2: **Early versus late normalization.** Each panel compares the performance, according to a specified performance measure, of a given resolution enhancement method when tested on LC Hi-C data subjected to early normalization (left two columns), and early normalization (x-axis) versus late normalization (y-axis)(right 2 columns). **a.** Structural similarity (SSIM) metric comparison. **b.** Peak signal-to-noise ratio (PSNR) comparison. **c.** Pearson's correlation coefficient (PCC) comparison. **d.** Spearman's correlation coefficient (PCC) comparison. **e.** Topologically Associating Domains (TAD) F1 comparison. **f.** Loop F1 comparison. **g.** Mean rank comparison.

This experiment yields the expected result: because the models are trained from data that employs early normalization, we observe much better performance on test data that uses this same early normalization approach (Figure S2). Two points are worth noting here. First, in practice, if you only collect LC Hi-C data, then the early normalization approach is impossible. Second, all previous methods have reported test set results that employ the late normalization approach [33, 41, 74, 21]. By doing normalization from the HC data before generating the corresponding LC data, these methods improperly leak information from the training set to the test set.

To address this problem, we propose a consistent analysis pipeline for normalized Hi-C (**Fig. S1b**). During training, the raw Hi-C data is first downsampled to generate raw LC Hi-C. Both raw Hi-C and raw LC Hi-C are then normalized separately, resulting in normalized Hi-C and normalized LC Hi-C, which serve as the training target and testing target, respectively. Critically, when a new LC Hi-C matrix is generated, it can be normalized without reference to any HC Hi-C data and then directly input to the model.

To empirically test whether the choice of normalization procedure impacts performance, we retrained a variety of methods (HiCARN1, HiCARN2, HiCNN, HiCSR) using data generated by the old (early normalization) and new (late normalization) pipelines. For comparison with other methods' pre-trained models, we adhered to the same training protocols as they did. In particular, the training data includes only training chromosomes (excluding chromosomes 4, 5, 11, and 14) from the GM12878 Hi-C data [71], and the HC target is clamped to [0, 255] and then normalized to [0, 1]. After retraining, we benchmarked using the normalized Hi-C from the same testing set as we used for raw Hi-C resolution enhancement, as described in Methods. The testing input is generated by first downsampling the HC raw Hi-C data and then applying KR normalization. The enhanced normalized Hi-C is then compared to the KR normalized HC Hi-C to evaluate the performance of different methods.

The overall improvement from the late normalization setting compared with the early normalization setting (in right two columns of **Fig. S2**) verifies our claim that the early normalization approach yields worse performance in practice. In general, the late-normalized models yielded better mean ranks than the corresponding early-normalized models. The only exception is HiCNN, which exhibited the opposite trend. This reduction may be because the input from normalized LC Hi-C increased the difficulty of convergence for a simple CNN network.
